# Rational engineering of sdAb-based CAR T cells targeting BCMA enhances antitumor efficacy and persistence in Multiple Myeloma

**DOI:** 10.64898/2026.06.02.729637

**Authors:** Vianca Ibarra-García, Maria E. Calleja-Cervantes, Federica Rochira, Iñigo Azagra, Patxi San Martin-Uriz, Paula Aguirre-Ruiz, Saray Rodriguez-Diaz, Rebeca Martinez-Turrillas, Elena Iglesias, Inés Ibañez-Sala, Eva Molina, Franco Bernasconi-Bisio, Paula Rodriguez-Marquez, Leonor Puchades-Carrasco, Nuria Gomez-Cebrian, Juan J. Lasarte, Mikel Hernaez, Ana Alfonso-Pierola, Esteban Tamariz, Paula Rodriguez-Otero, Jesus San Miguel, Juan R. Rodriguez-Madoz, Antonio Pineda-Lucena, Felipe Prosper

## Abstract

BCMA-directed CAR T therapies have transformed the treatment of relapsed/refractory multiple myeloma (MM), yet durable responses remain limited by insufficient persistence and functional exhaustion. The contribution of antigen-binding domain properties to CAR T cell performance is not fully defined. Here, we report the rational discovery and engineering of BCMA-targeting single-domain antibodies (sdAbs) and their incorporation into second-generation CAR T cells. We generated sdAbs recognizing distinct BCMA epitopes with diverse binding kinetics. Comprehensive biophysical and functional characterization identified sdAb5 as a lead candidate with balanced binding kinetics and a non-overlapping epitope relative to clinically approved constructs. sdAb-based CAR T cells exhibited potent antigen-specific cytotoxicity, minimal tonic signaling, and preserved IL-2 production without excessive inflammatory activation. In xenograft models, sdAb5-based CAR T cells induced durable tumor control, enhanced persistence, and improved responses to tumor rechallenge compared with ide-cel and cilta-cel constructs. Notably, this superior functionality correlated with balanced binding kinetics rather than maximal affinity. Single-cell transcriptomic analyses corroborated these findings, revealing a restrained inflammatory activation in sdAb5-based CAR T cells with preservation of transcriptional plasticity. Collectively, these findings support antigen-binding optimization as a key determinant of CAR T-cell durability and identify sdAb5 as a candidate for next-generation BCMA-directed therapies in MM.

## INTRODUCTION

Multiple myeloma (MM) remains an incurable plasma cell malignancy despite major therapeutic advances over the past decade^1,2^. Chimeric antigen receptor (CAR) T cell therapies targeting B-cell maturation antigen (BCMA) have produced unprecedented response rates in patients with relapsed or refractory disease^3,4^, leading to the clinical approval of idecabtagene vicleucel (ide-cel; Abecma) and ciltacabtagene autoleucel (cilta-cel; Carvykti)^5,6^. Nevertheless, durable remissions are limited to a subset of patients, and relapse remains frequent^7,8^. Mechanisms of failure include insufficient CAR T cell persistence, progressive functional exhaustion, and therapeutic resistance following prior BCMA-directed interventions^9,10^. As BCMA CAR T cells are increasingly deployed earlier in the treatment paradigm^5,6^, improving durability, functional fitness, and reusability has emerged as a central clinical challenge.

The antigen-binding domain of a CAR is a critical but often underappreciated determinant of therapeutic performance^11^. Most clinically approved CAR T cell products rely on single-chain variable fragments (scFv), which are susceptible to tonic signaling, aggregation, and structural instability^12–14^. These intrinsic properties may contribute to premature T cell dysfunction, limited persistence and exacerbated inflammatory toxicity. In these sense single-domain antibodies (sdAbs) offer several features that make them particularly attractive for CAR engineering. Their small size and monomeric nature confer superior stability and reduce the propensity for aggregation and tonic signaling^13,14^. sdAbs can access cryptic or membrane-proximal epitopes that may be sterically restricted to conventional antibodies, thereby expanding the targetable epitope landscape^15,16^. In addition, their modular architecture facilitates the generation of multivalent or multispecific CAR designs with precise spatial configuration. Notably, cilta-cel which incorporates two sdAbs in a biparatopic configuration, represents a distinct design paradigm with demonstrated remarkable clinical efficacy^4,9^. Collectively, these attributes position sdAbs as a versatile platform for next-generation CAR design, although how sdAb-intrinsic properties, including epitope selection, binding kinetics, and structural configuration, translate into functional CAR T cell outcomes remains poorly understood.

Beyond affinity, the kinetics and epitope specificity of CAR-antigen interactions critically shape T cell fate. Excessively stable binding may impair serial target engagement and promote exhaustion, whereas suboptimal interactions may limit efficacy^17^. Moreover, epitope overlap among BCMA-targeted therapies may facilitate resistance in the setting of sequential treatment^9,18^. Rational selection of antigen-binding domains with optimized kinetics and non-overlapping epitope engagement therefore represents a promising strategy to enhance both the durability and clinical versatility of CAR T cell therapies.

Recent advances in single-cell transcriptomics have further refined our understanding of CAR T cell function^19^ by revealing the heterogeneity and dynamic evolution of T cell states *in vivo*^20–24^. These studies indicate that durable responses are associated with balanced activation programs and preserved transcriptional plasticity, whereas excessive inflammatory signaling drives differentiation and dysfunction. However, how CAR design, particularly the antigen-binding domain, shapes these transcriptional trajectories remain incompletely understood.

In this study, we report on the rational discovery and engineering of BCMA-specific sdAbs and their incorporation into second-generation CAR T cells. Using llama immunization coupled with phage display, we generated a diverse panel of high affinity sdAbs recognizing distinct BCMA epitopes. We systematically evaluated their biophysical properties, signaling behavior, and antitumor activity, benchmarking lead candidates against ide-cel and cilta-cel constructs. Moreover, using *in vivo* xenograft models and longitudinal single-cell transcriptomic profiling, we interrogated the cellular programs underlying persistence, exhaustion, and response to tumor rechallenge.

## RESULTS

### Generation of a diverse panel of high-affinity BCMA-targeting single-domain antibodies

To generate novel single-domain antibodies (sdAbs) targeting B-cell maturation antigen (BCMA), we immunized a llama with recombinant human BCMA. Immunization elicited a robust humoral response, with strong titers of anti-BCMA antibodies, detected by ELISA, as early as the first injection. By the third dose, titers exceeded 1:10,000, and polyclonal serum displayed an EC₅₀ of 49.3 nM (Figure S1A). Following immunization, a phage display library containing ∼10^10^ unique clones was constructed and subjected to four rounds of panning under increasingly stringent conditions (Figure S1B). The output/input phage ratio increased by three orders of magnitude, and polyclonal phage ELISA confirmed enrichment for BCMA-binding clones, with a signal-to-noise ratio >50:1 in the final round (Figure S1C). No binding to unrelated His-tagged proteins or bovine serum albumin (BSA) was observed, confirming target specificity. Sequence analysis of selected clones revealed extensive diversity, with 30% unique CDR3 and 50% unique CDR1/CDR2 regions, resulting in 23 unique sdAbs clustered into 6 different families (Figure S1D). Thus, these results provide us with different BCMA-specific sdAbs that were further characterized.

### Distinct binding kinetics and epitope engagement define functionally diverse BCMA sdAbs

Five representatives, each one from a different family (sdAb5, sdAb10, sdAb15, sdAb18, and sdAb22), were selected for detailed characterization (Figure S1E). Recombinant expression of selected sdAbs in *E. coli* yielded 1-5 mg/L of protein, which were further purified, via immobilized metal affinity chromatography (IMAC) and ion-exchange chromatography (IEC), reaching >95% homogeneity as confirmed by SDS-PAGE (Figure S1F). Purified sdAbs demonstrated strong binding to human BCMA in ELISA, with sigmoidal dose-response curves and EC₅₀ values in the low-nanomolar range (Figure S1G). Surface plasmon resonance (SPR) revealed a different range of affinities driven by distinct kinetic profiles (association and dissociation rates) (Figure 1A, Table S1). sdAb10 showed the highest affinity (K_D_ = 1.75×10^−10^ M) resulting from a rapid association (k_on_ = 6.88×10^6^ M^−1^s^−1^) and a slow dissociation rate (k_off_ = 1.07×10^−3^ s^−1^). In contrast, sdAb15 exhibited the weakest binding (K_D_ = 1.16×10^−7^ M), with low association and high dissociation rates (k_on_ = 1.17×10^5^ M^−1^s^−1^; k_off_ = 1.36×10^−2^ s^−1^). sdAb5 displayed high affinity (K_D_ = 8.62×10^−10^ M) with balanced kinetic parameters (k_on_ = 3.07×10^6^ M^−1^s^−1^; k_off_ = 2.40×10^−3^ s^−1^) (Table S1). Epitope binning by bio-layer interferometry (BLI) demonstrated that sdAb5 and sdAb10 recognized non-overlapping epitopes on human BCMA, whereas sdAb15, sdAb18, and sdAb22 bound to regions partially or fully overlapping with sdAb5 and/or sdAb10 (Figure 1B-C). Structural modeling, using the Molecular Operating Environment (MOE) platform, was consistent with these findings, suggesting distinct binding interfaces for each sdAb on the BCMA extracellular domain (Figure 1D). Together, these data establish a panel of high affinity sdAbs suitable for CAR engineering.

**Figure 1.**
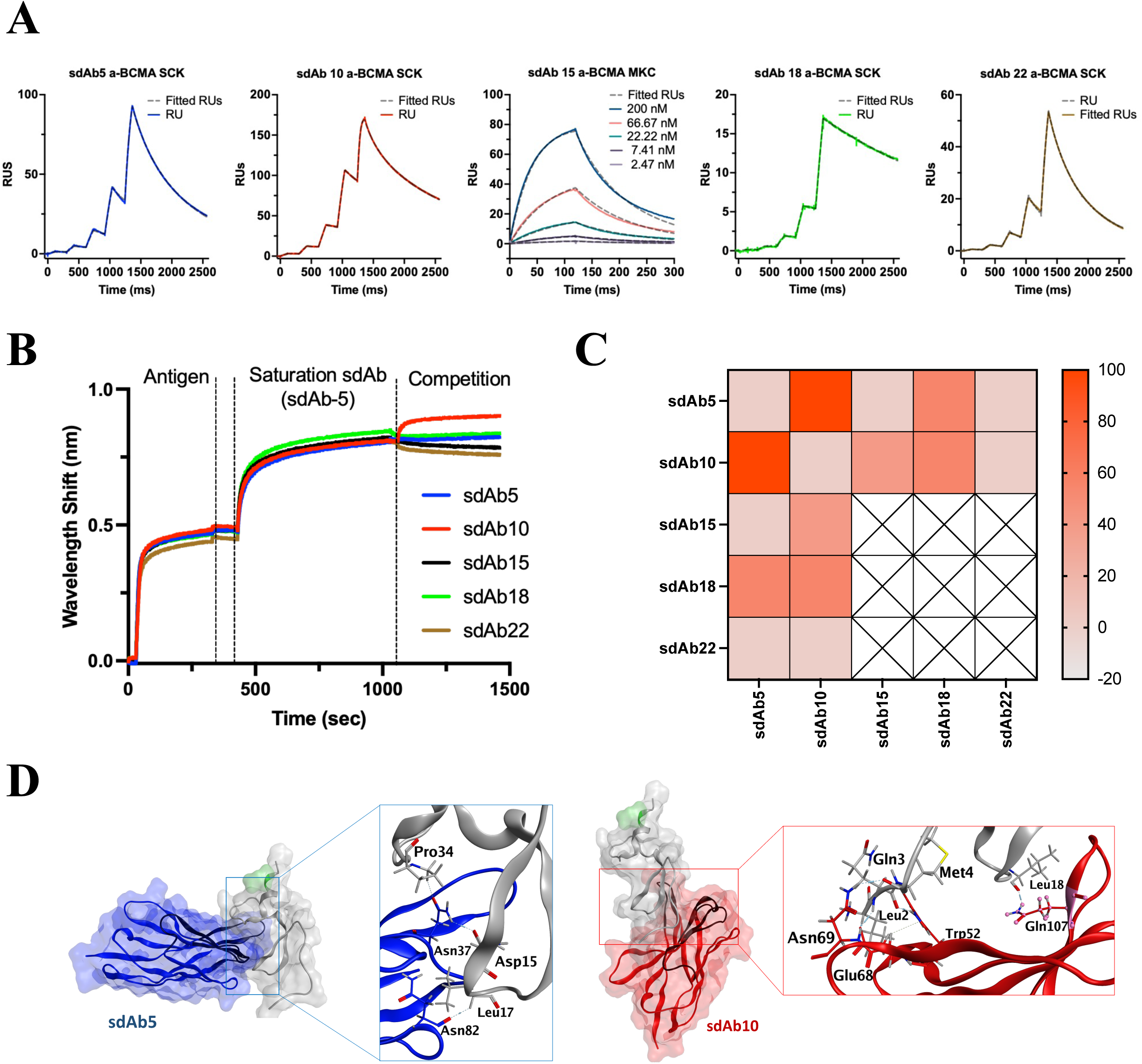
Biophysical and structural characterization of BCMA-binding single-domain antibodies (sdAbs). **(A)** Surface plasmon resonance (SPR) sensorgrams showing kinetic binding profiles of each sdAb, including single-cycle kinetics (SCK) for sdAb5, sdAb10, sdAb18, sdAb22 and multi-cycle kinetics (MCK) for sdAb15. **(B)** Bio-layer interferometry (BLI)-based competition assay to assess pairwise binding interference. **(C)** Epitope binning heatmap derived from BLI data. Dark red indicates complete competition (overlapping epitope); light red denotes partial competition or distinct binding sites. **(D)** Computational docking of sdAb5 and sdAb10 on the BCMA ectodomain highlighting predicted interaction surfaces and potential competitive overlap.

### sdAb-based CARs exhibit antigen-specific activation with minimal tonic signaling

To assess the functionality and specificity of selected sdAbs as recognition moieties of a CAR, several constructs were designed by incorporating each sdAb into a second generation 4-1BB CAR structure fused to a truncated human EGF receptor (EGFRt) as surrogate marker (Figure S2A). Jurkat triple parameter reporter (TPR) cells, used to assess tonic signal and activation, were transduced with selected sdAb-CAR T cells and proper CAR expression was confirmed by EGFRt co-expression (Figure S2B). Tonic signaling was evaluated by combining basal Jurkat TPR activity (in the absence of antigen) with computational *in silico* modeling of positively charged patches (PCPs) (Figure S2C-F). Almost all sdAb-based CAR constructs presented minimal reporter activation and PCP values <40 (previously defined threshold^13,14^), only sdAb18 displayed elevated background activation and increased PCP value (>40) (Figure S2G). Upon co-culture with BCMA^+^ MM1S cells, again all sdAb-based constructs but sdAb18, induced strong NFAT, NF-κB, and AP-1 reporter activation (Figure S2D-F), confirming antigen-dependent signaling. Given the association of tonic signaling with T cell dysfunction, sdAb18 was excluded from further studies in human T cells.

Next, we performed a deep phenotypic and functional characterization of CAR T cells, generated with the selected sdAb constructs. As control we designed CAR T cells with the same binder domain of biparatopic sdAb-based CAR T product cilta-cel and a humanized version of the scFv from ide-cel (C11D5.3h). All CAR T products showed high transduction rates, with >50% EGFRt^+^ cells, together with an efficient *in vitro* expansion (Figure 2A and S3A). Moreover, no significant differences in the distribution of CD4⁺ and CD8⁺ T cell populations and memory/effector subset composition were observed across constructs (Figure 2B and S3B). Importantly, all CAR T cells maintained low expression of exhaustion markers (PD-1, TIM-3, LAG-3, TIGIT), presenting modest upregulation of activation markers (HLA-DR, CD137, CD69, ICOS), again without significant differences between products (Figure 2C and S3C). Cytotoxicity assays confirmed potent and antigen-specific activity of sdAb-based CAR T cells, with all novel sdAb-based constructs presenting similar lytic capacity than cilta-cel and C11D5.3h (Figure 2D and S3D-E). Moreover, cytotoxic activity correlated with BCMA density across myeloma lines (MM1S, U266, H929, MOLP8) underscoring dependence on antigen expression (Figure S3F-G). After antigen stimulation, sdAb-based CAR T cells secreted higher levels of IL-2 compared with cilta-cel and C11D5.3h (Figure 2E), suggesting enhanced proliferation and persistence potential. In contrast, IFN-γ, TNF-α, and granzyme B production were comparable across all CAR T cells, reflecting strong effector function. Extended cytokine profiling by multiplex analysis showed very similar profiles, with only IL-8 presenting modest increase in sdAb-based CAR T cells relative to cilta-cel (Figure S4). Overall, our novel sdAb-based CAR T cells displayed effector function without evidence of hyperactivation in *in vitro* studies.

**Figure 2.**
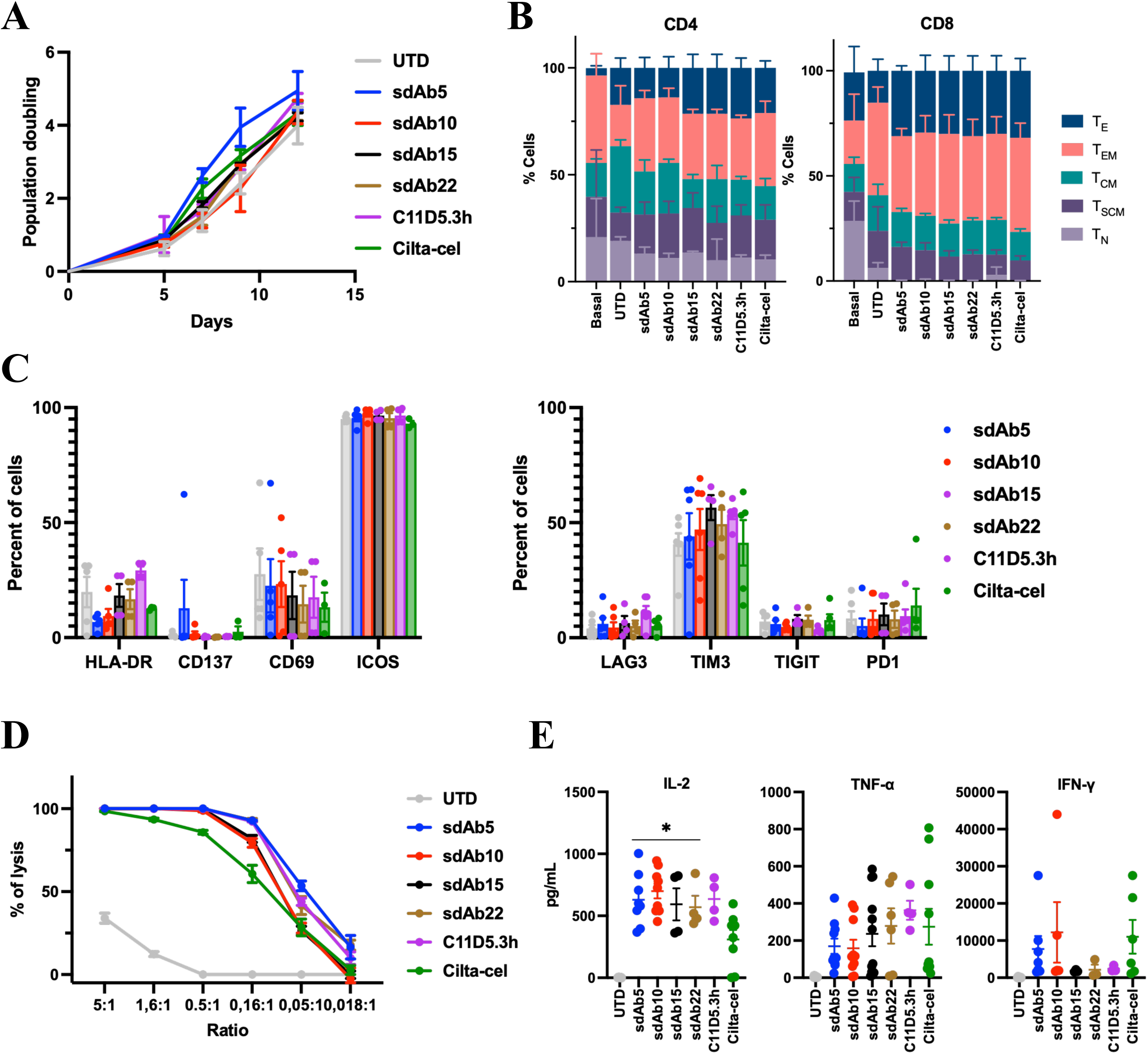
*In vitro* functional profiling of BCMA-directed sdAb-based CAR T cells. **(A)** Expansion kinetics of sdAb-based CAR T cells during manufacturing. **(B)** Differentiation profile of CD4⁺ (left) and CD8⁺ (right) CAR T cell subsets, showing distribution among naïve (T_N_), stem cell memory (T_SCM_), central memory (T_CM_), effector memory (T_EM_), and effector (T_E_) compartments. **(C)** Expression of activation (HLA-DR, CD137, CD69, ICOS) and exhaustion (LAG3, TIM3, TIGIT, PD1) markers on CD8⁺ CAR T cells following 24-hour co-culture with MM1S target cells. **(D)** *In vitro* cytotoxicity against MM1S-Luc⁺ cells at multiple effector-to-target (E:T) ratios. **(E)** Cytokine secretion profile (IL-2, TNF-α and FN-γ) in co-culture supernatants, quantified by Luminex multiplex assay (broader multiplex profiling is shown in Figure S4.).

### sdAb-based CAR T cells mediate robust long-term *in vivo* antitumor activity

Next, we further assessed *in vivo* antitumoral efficacy of developed sdAb-based CAR T cells in a xenogeneic MM model in NSG mice (Figure S5A). We observed that a single intravenous administration of 10^5^ CAR T cells in MM1S-Luc⁺ tumor bearing mice (a suboptimal dose to enable detection of potential differences), significantly prolonged survival of treated animals compared with control groups (PBS and UTD) (Figure S5B). In particular, sdAb5-based CAR T cells presented the highest efficacy, with 50% of mice surviving beyond day 162, followed by sdAb10-, sdAb15-, and sdAb22-based CAR T products (Figure S5B). These outcomes were broadly consistent with the biophysical properties of the corresponding sdAbs (Table S1), with sdAb5– and sdAb10-based CAR T cells showing superior *in vivo* activity compared with constructs incorporating lower-affinity binders. However, the observed differences in antitumoral efficacy were not strictly proportional to binding affinity, indicating that additional parameters beyond K_D_ contribute to optimal CAR T cell function *in vivo*. Collectively, these findings position sdAb5– and sdAb10-based CAR T cells as lead candidates combining strict antigen specificity, favorable cytokine profiles, and durable *in vivo* efficacy, and providing the rationale to advance sdAb5– and sdAb10-based CAR T cells for further characterization.

A direct comparison of the two lead sdAb-based CAR T cell products with C11D5.3h revealed a similar antitumoral efficacy of sdAb10-based CAR T cells and a significantly prolonged survival of animals treated with sdAb5-based CAR T cells, with 50% of the animals alive at day 150 post-infusion, highlighting the improved antitumoral response of our novel CAR T product (Figure 3A). Moreover, in a head-to-head comparison with cilta-cel, sdAb5-based CAR T cells achieved a comparable efficacy (Figure 3B), with a concomitant reduction of the tumor burden measured by bioluminescence (Figure 3C). In addition, sdAb5-based CAR T cells achieved a higher expansion at day 7 post administration within the bone marrow, inducing a more rapid tumor clearance (Figure 3D). These findings suggest that sdAb5-based CAR T cells facilitate superior early-phase expansion and persistence within the tumor microenvironment, contributing to both immediate and durable anti-tumor efficacy. We further hypothesized that combining sdAb5 and sdAb10 into a single biparatopic CAR construct, analogous to the cilta-cel design, might further enhance antitumoral activity by increasing avidity and epitope coverage (Figure S5C). Although this biparatopic construct displayed markedly increased affinity, it did not improve *in vivo* efficacy and was associated with reduced survival compared with sdAb5-based CAR T cells alone (Figure S5D). While high-affinity binding is often considered advantageous, excessively stable interactions may impair serial target engagement, limit synapse turnover, or promote dysfunctional signaling.

**Figure 3.**
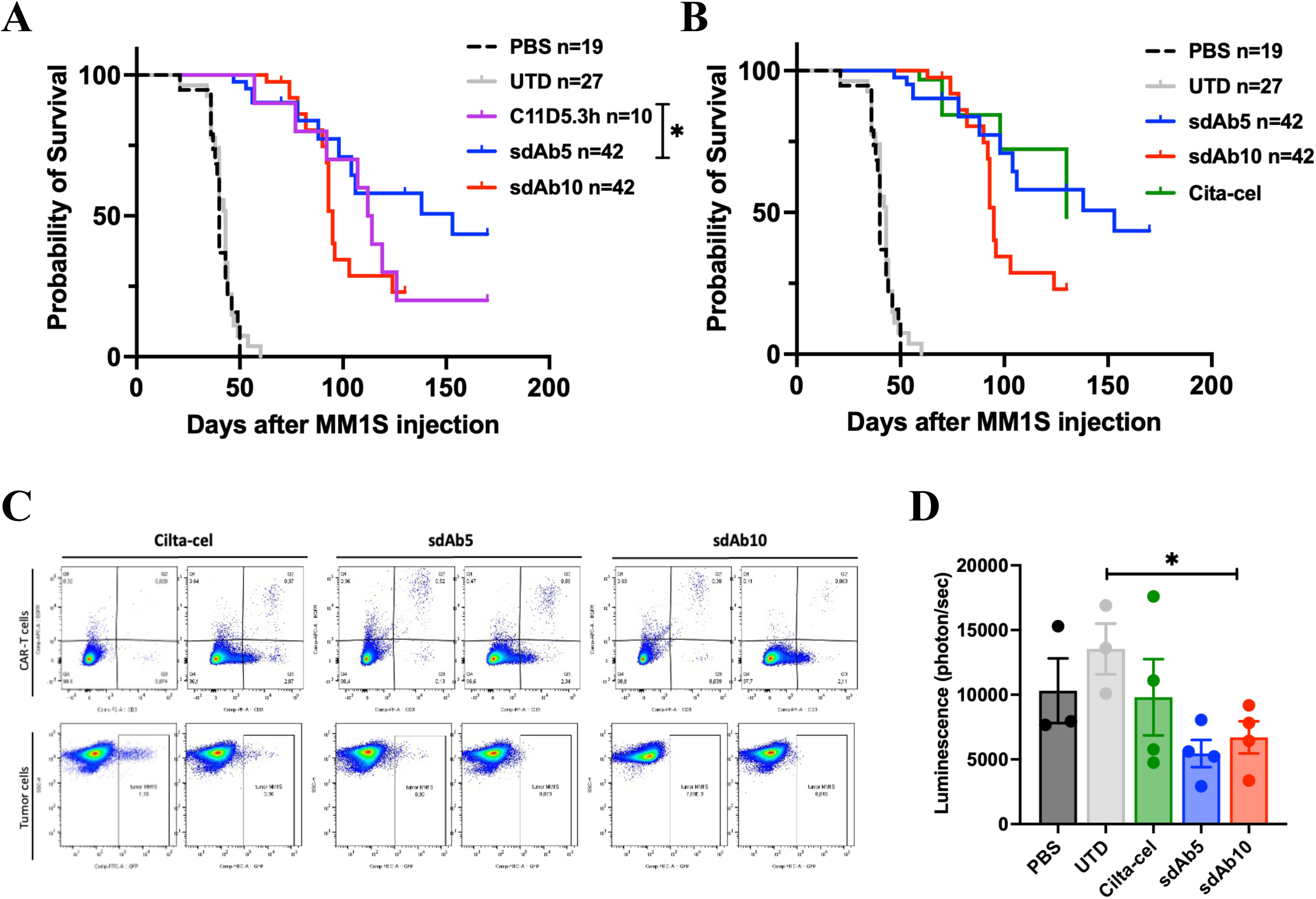
*In vivo* efficacy and persistence of sdAb-based CAR T cells in a xenograft multiple myeloma model. **(A)** Kaplan-Meier survival curves of NSG mice intravenously injected with MM1S-Luc⁺ tumor cells and treated 14 days later with CAR T cells incorporating sdAb5, sdAb10, or the humanized C11D5.3h binder. Untreated (PBS) and untransduced (UTD) T cell groups served as controls (n=5-10 per group). **(B)** Comparative survival analysis of selected sdAb-based CAR constructs versus Cilta-cel. **(C)** Flow cytometry plots showing CAR T cell persistence (CD3⁺EGFR⁺) and residual MM1S tumor burden (GFP⁺) in the bone marrow on day 7 post-infusion. **(D)** Quantification of bioluminescent tumor signal (photons/sec) in mice treated with sdAb5, sdAb10, or Cilta-cel CAR T cells on day 7 post-infusion.

### sdAb5-based CAR T cells mediate superior and durable protection against tumor rechallenge

Then, we further evaluate the long-term persistence and functionality of our selected sdAb-based CAR T cells assessing their efficacy against a tumor rechallenge, as a model of tumor relapse. Thus, mice treated with sdAb5-, sdAb10-based CAR T cell or cilta-cel were re-infused with MM1S-Luc⁺ cells at day 70, and disease progression was monitored by bioluminescence and survival analysis (Figure 4A). Despite some residual disease was observed before rechallenge in some mice (Figure 4B), animals previously treated with sdAb5– and sdAb10-based CAR T cells were more effective controlling tumor growth than those treated with cilta-cel, effect that was translated into a superior survival rate (Figure 4C). Moreover, flow cytometry analysis confirmed higher CAR T cell frequencies (CD3⁺EGFR⁺ cells) in bone marrow and spleen of sdAb5-based CAR T cells treated mice at day 7 and 14 after rechallenge, indicating robust *in vivo* re-expansion capacity (Figure 4D). These results demonstrate not only the long-term persistence of sdAb5-based CAR T cells but also their ability to achieve effective long-term responses, a critical property for preventing relapse, providing strong evidence of potential durable therapeutic responses of sdAb-based CAR T constructs in MM.

**Figure 4.**
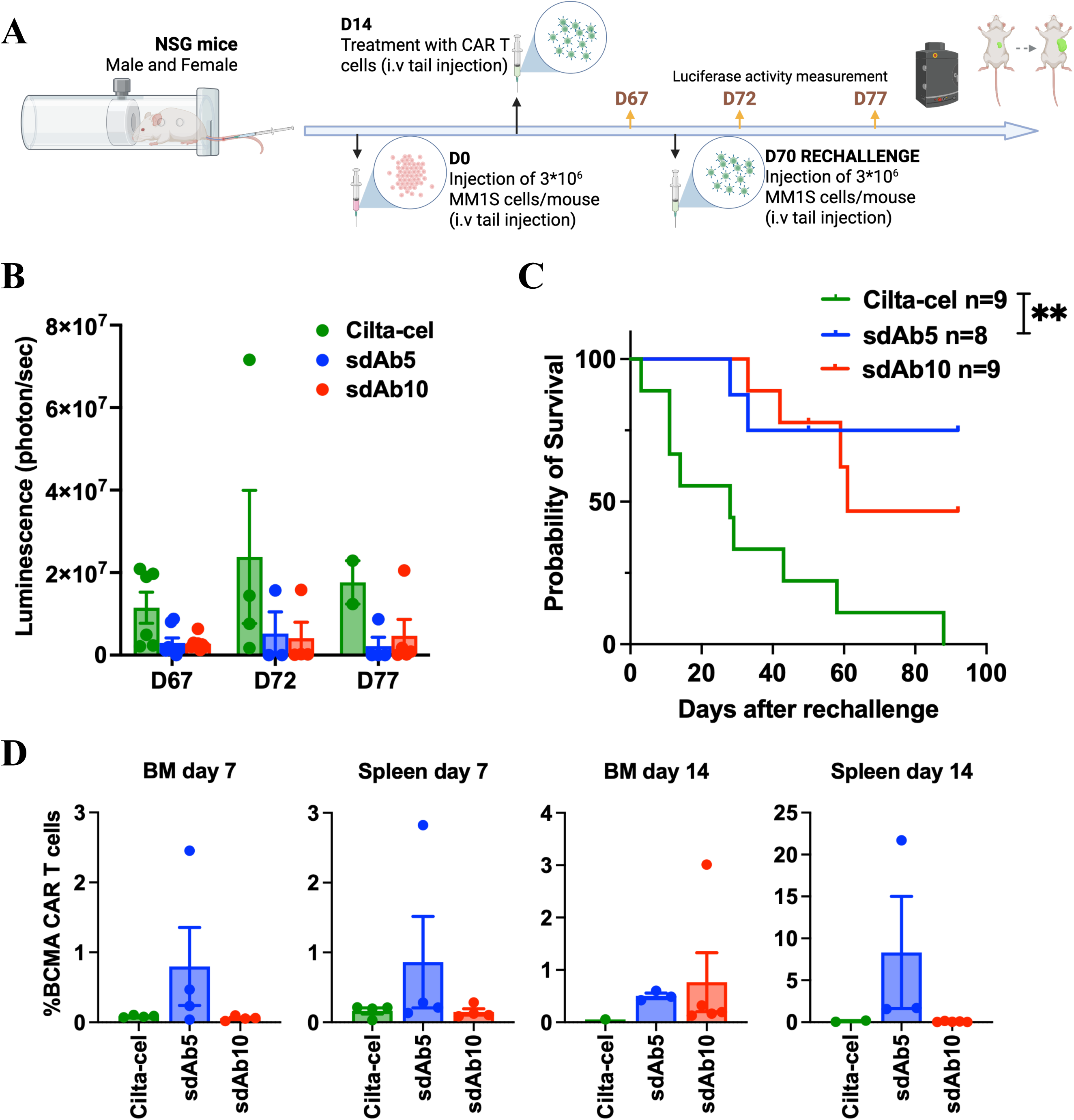
sdAb-based CAR T cells demonstrate durable anti-tumor responses and persistence following tumor rechallenge. **(A)** Experimental timeline of tumor rechallenge model. NSG mice bearing MM1S-Luc⁺ tumors were treated with sdAb-based CAR T cells (sdAb5, sdAb10, or Cilta-cel). After tumor clearance and long-term monitoring, mice were rechallenged intravenously with MM1S-Luc⁺ cells on day 70. **(B)** Tumor burden was quantified by *in vivo* bioluminescence imaging at days 67 (before rechallenge), 72 and 77 (day 2 and 7 post-rechallenge). **(C)** Kaplan-Meier survival analysis following rechallenge, showing enhanced protection and long-term survival in mice previously treated with sdAb5-based CAR T cells. **(D)** Flow cytometric quantification of CAR T cells (CD3⁺EGFR⁺) in bone marrow and spleen on days 7 and 14 post-rechallenge.

### sdAb5 engages a non-overlapping BCMA epitope with distinct properties compared to ide-cel and cilta-cel binders

To decipher the underlying mechanisms driving differences in CAR T cells functionality, we benchmarked our sdAbs against the sdAbs present in cilta-cel (cilta-1 and cilta-2), as well as with the scFv from ide-cel (C11D5.3) and its humanized version (C11D5.3h) used in this study. Binding kinetics were defined using multi– and single-cycle SPR. Interestingly, we observed that the affinity of sdAb5 was in range with the biparatopic region of cilta-cel (K_D_ = 8.62×10^−10^ M vs K_D_ = 2.79×10^−10^ M), supporting the similar antitumoral efficacy observed between these constructs. On the other hand, sdAb10 matched the affinity of C11D5.3 (K_D_ = 1.75×10^−10^ M vs K_D_ = 1.05×10^−10^ M), and C11D5.3h displayed the strongest affinity (K_D_ = 1.05×10^−11^ M) due to an exceptionally slow off-rate (Figure 5A-F, S6A-B and Table S1). Further characterization of epitope mapping, evaluated by BLI and structural modeling, showed that sdAb5 engaged an epitope with no competition with any of the two sdAb components of cilta-cel (Figure 5G and S6C-D). By contrast, sdAb10 exhibited partial competition with both cilta-1 and cilta-2 (Figure 5H and S6C-D), and docking analysis revealed overlapping with cilta-2, suggesting adjacent or shared epitope engagement (Figure S6E). Competition assays confirmed that biparatopic cilta-cel displaced sdAb10 but not sdAb5 (Figure 5I). Notably, C11D5.3h blocked binding of all constructs, consistent with its broad epitope footprint (Figure S6F-G). Collectively, these findings identify sdAb5 as a BCMA binder with a distinct epitope and dynamic binding profile compared with the recognition domains used in currently approved CAR T cell therapies. The non-overlapping epitope engagement of sdAb5, together with its favorable functional and persistence characteristics, suggests that sdAb5-based CAR T cells may retain activity in settings where tumor escape is driven by epitope masking or selective pressure from prior BCMA-targeted therapies.

**Figure 5.**
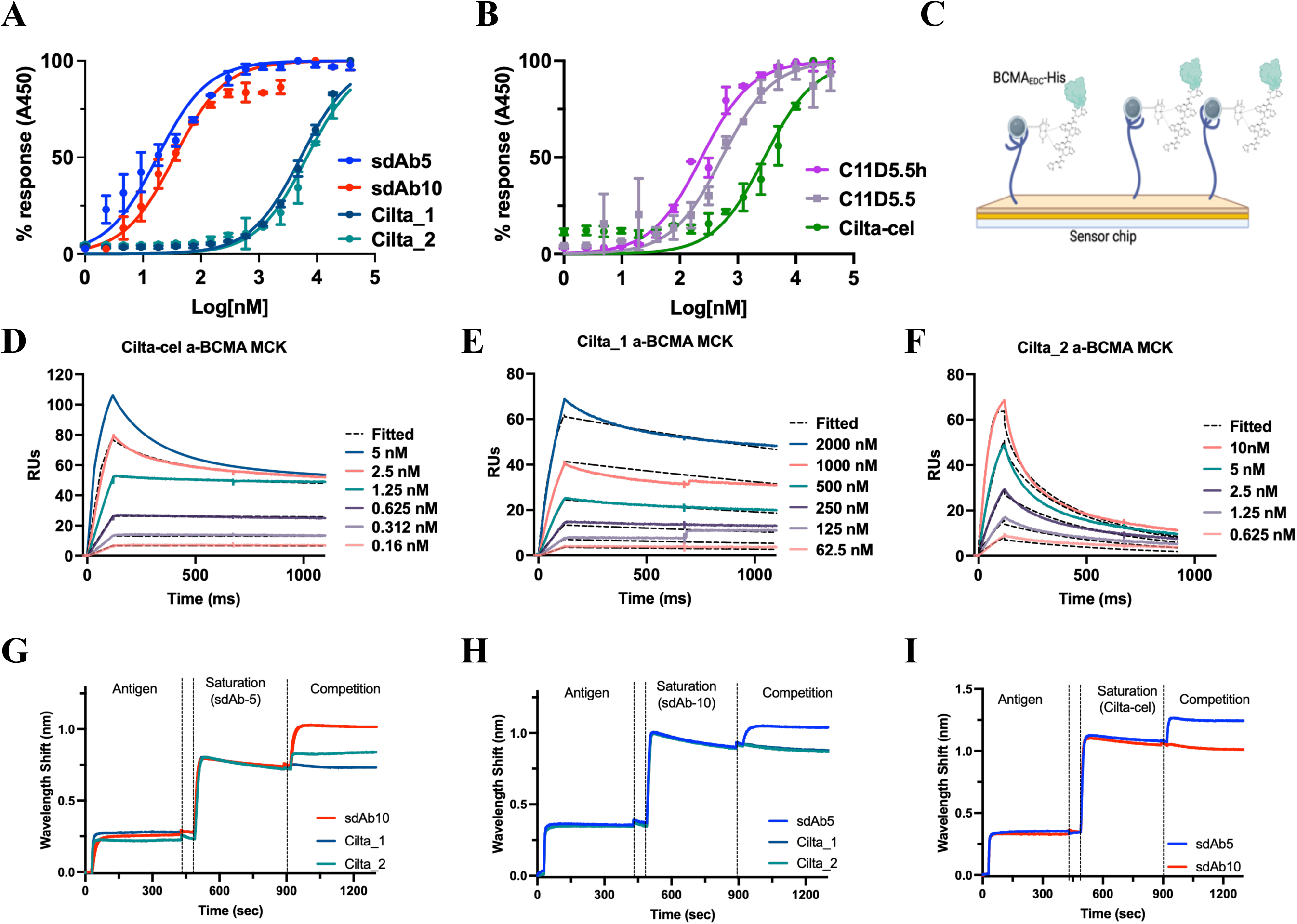
Comparative biophysical analysis of sdAb5 and sdAb10 binders. **(A)** ELISA-based binding curves showing the affinity of sdAb5, sdAb10, and Cilta-cel monomers (Cilta_1, Cilta_2). **(B)** ELISA-based binding curves showing the affinity of full biparatopic Cilta-cel and the scFv C11D5.3/C11D5.3h used in ide-cel. **(C)** Schematic of the surface plasmon resonance (SPR) assay setup, with His-tagged BCMA immobilized on the chip. **(D-F)** Multi-cycle kinetic (MCK) SPR analysis of biparatopic Cilta-cel (D), Cilta_1 (E) and Cilta_2 (F) revealing binding kinetics and affinity constants. **(G-I)** Bio-layer interferometry (BLI)-based competition assays assessing epitope overlap: (G) sdAb5 vs. sdAb10 and Cilta-cel components, (H) sdAb10 vs. sdAb5 and Cilta-cel components, and (I) biparatopic Cilta-cel vs. sdAb5 and sdAb10, demonstrating distinct or overlapping binding profiles.

### Single-cell transcriptomics reveal restrained inflammatory activation programs of sdAb-based CAR T cells

To investigate the transcriptional programs underlying the improved antitumor activity of sdAb5-based CAR T cells, we performed single-cell RNA sequencing of almost 25,000 CAR T cells isolated from BM of control mice and mice treated with sdAb5 (9,581 cells), sdAb10 (7,703 cells) and Cilta-cel (7,204 cells) during the primary response phase (day 7), prior to tumor rechallenge (day 70), and following tumor rechallenge (day 77). Unsupervised clustering and annotation identified 10 transcriptionally distinct T-cell states spanning both CD4 and CD8 lineages, including naïve/central memory-like populations, cytotoxic T lymphocytes (CTL), exhausted cells (Tex), Th1, Th2, regulatory T cells (Treg), and cycling effector subsets (Figure 6A-B and S7A). Although cells from all CAR T constructs were represented across clusters, their relative distribution varied markedly across time points. While contributions were broadly similar during the early response phase, several clusters at later stages became strongly enriched or nearly exclusive for specific CAR T products (Figure 6C and S7B-C).

**Figure 6.**
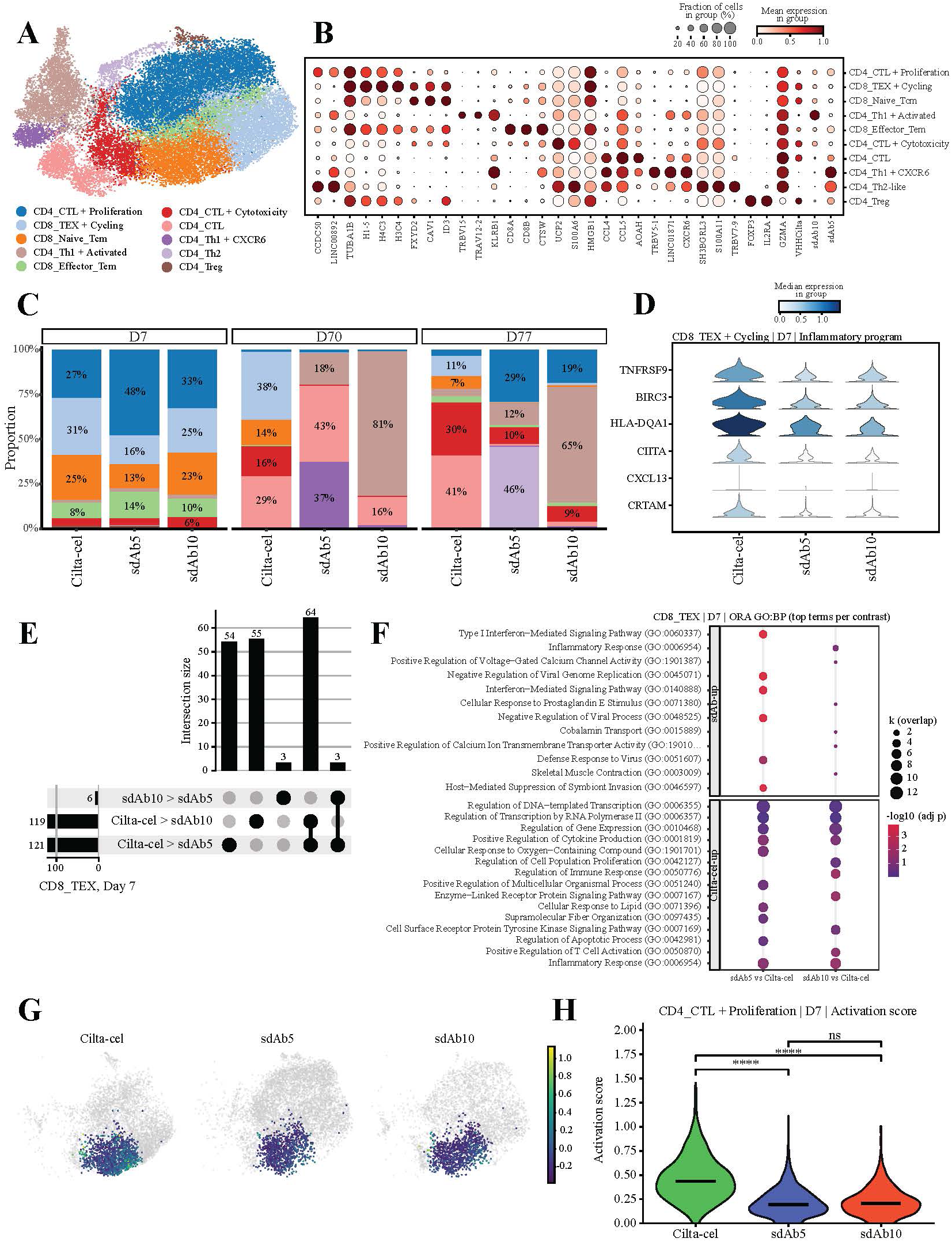
Mechanism of sdAb-based CAR T treatment in primary efficacy and after tumor rechallenge. **(A)** UMAP of CAR-positive T cells colored by T-cell states. **(B)** Dotplot summarizing representative marker genes across the 10 transcriptional cell states. **(C)** Stacked composition plots showing the distribution of T-cell states across treatments within each consolidated sampling window. **(D)** Violin plots showing expression of selected activation– and antigen-presentation-associated genes in CD8_TEX cells across treatments at D7. **(E)** UpSet plot summarizing overlap of globally downregulated genes in CD8_TEX cells at D7 across pairwise treatment comparisons. **(F)** GO biological process enrichment analysis of genes differentially regulated in sdAb5 or sdAb10 relative to Cilta-cel in CD8_TEX cells. **(G)** UMAP showing the distribution of the exhaustion score across treatments at D7 in CD8_Naive_Tcm cells. **(H)** Violin plots comparing activation score across treatments at D7 in CD4_CTL cells.

During the primary efficacy phase (day 7), all CAR T cell products displayed robust activation, characterized by high proliferative activity (G2M/S) and enrichment of cycling cytotoxic transcriptional programs (Figure S8A). Across constructs, CAR T compartment was dominated by three major populations, CD4_CTL + Proliferation, CD8_TEX, and CD8_Naive_Tcm, together representing more than 75% of recovered cells (Figure 6C and Table S2). This shared early landscape indicates that all products initiate a similar primary effector response following antigen engagement. Despite these similarities, subtle but consistent transcriptional differences emerged between products. In particular, sdAb5-based CAR T cells showed a shift toward CD4_CTL + Proliferation populations, whereas cilta-cel was relatively enriched in CD8_TEX cells (Figure 6C and S8B). Further differential gene expression analysis within the CD8_TEX compartment revealed stronger inflammatory and activation-associated transcriptional programs in cilta-cel, driven by preferential upregulation of inflammatory-response genes (Figure 6D-E). Consistent with this observation, over-representation analysis highlighted enrichment of cytokine signaling, immune activation, and inflammatory pathways in cilta-cel (Figure 6F). Differences were also observed within the CD8_Naive_Tcm compartment, where cilta-cel showed stronger enrichment of regulatory and signaling processes (Figure S9). Projection of an early T-cell receptor activation signature further revealed a preferential enrichment of immediate activation genes (including CCL3, NR4A1, CRTAM, IL2RA, and DUSP4) in cilta-cel (Figure 6G). A similar trend was observed within the CD4_CTL + Proliferation population, in which cilta-cel exhibited higher activation scores driven by genes associated with early activation and co-stimulation, including TNFRSF9, ICOS, IL2RA, and CXCL13 (Figure 6H). Gene ontology enrichment analysis further confirmed stronger enrichment of transcriptional regulation and inflammatory signaling pathways in this population (Figure S9).

Collectively, these data indicate that although all CAR T cell products undergo rapid early activation following antigen encounter, sdAb-based CAR T cells adopt a distinct transcriptional landscape characterized by restrained inflammatory signaling and increased representation of proliferative CD4 cytotoxic populations. Compared with cilta-cel, this more balanced effector program may reflect reduced early overstimulation, a feature that could contribute to the enhanced persistence and long-term functional adaptability observed with sdAb-based constructs.

### sdAb-based CAR T cells exhibit enhanced transcriptional plasticity during long-term persistence and tumor rechallenge

At the long-term pre-rechallenge time point (day 70), differences between CAR T cell products became more pronounced, indicating divergent long-term differentiation trajectories, with sdAb-based CAR T cells showing preferential persistence of non-proliferative oligoclonal CD4 helper-like transcriptional states, without evidence of large clonal expansions (Figure 6C, 7A-B and S10). In particular, sdAb5-based CAR T cells were enriched in a CXCR6-expressing CD4_Th1 subset, whereas sdAb10-based CAR T cells favored more broadly activated CD4_Th1 programs. In contrast, cilta-cel CAR T cells remained largely dominated by more polyclonal CD8_TEX populations, consistent with a more differentiated and exhaustion-skewed phenotype (Figure 7A and S8B). Interestingly, following tumor rechallenge, CAR T cell transcriptional responses markedly diverged across constructs, with sdAb-based products showing a robust emergence of CD4_CTL + Proliferation populations, indicating renewed engagement of cytotoxic effector programs. Notably, sdAb5 CAR T cells underwent substantial transcriptional remodeling towards the emergence of a CD4_Th2-like program, whereas sdAb10 preferentially expanded activated CD4_Th1 populations (Figures 7A and S8B). This specific sdAb5-associated CD4_Th2 program was marked by upregulation of canonical Th2 polarization genes (GATA3, IL-13, IL-5) and enrichment of immune signaling pathways (Figures 7C-D), suggesting a helper-oriented functional program that may support sustained effector activity during repeated antigen encounters. In contrast, cilta-cel CAR T cells displayed limited transcriptional remodeling following rechallenge, with expansions largely restricted to CD4_CTL states already prevalent prior to rechallenge, suggesting a more constrained secondary response. Consistently, pseudobulk analyses within the CD4_CTL compartment revealed that cilta-cel cells maintained stronger activation and interferon-associated transcriptional programs (Figure 7E). Moreover, although prolonged antigen exposure ultimately led to induction of exhaustion-associated transcriptional programs (PDCD1, TOX, TOX2, LAG3, HAVCR2, TIGIT, and BATF) broadly distributed across cell states, these signatures were significatively more prominent on cilta-cel (Figure 7F-G and S11).

**Figure 7.**
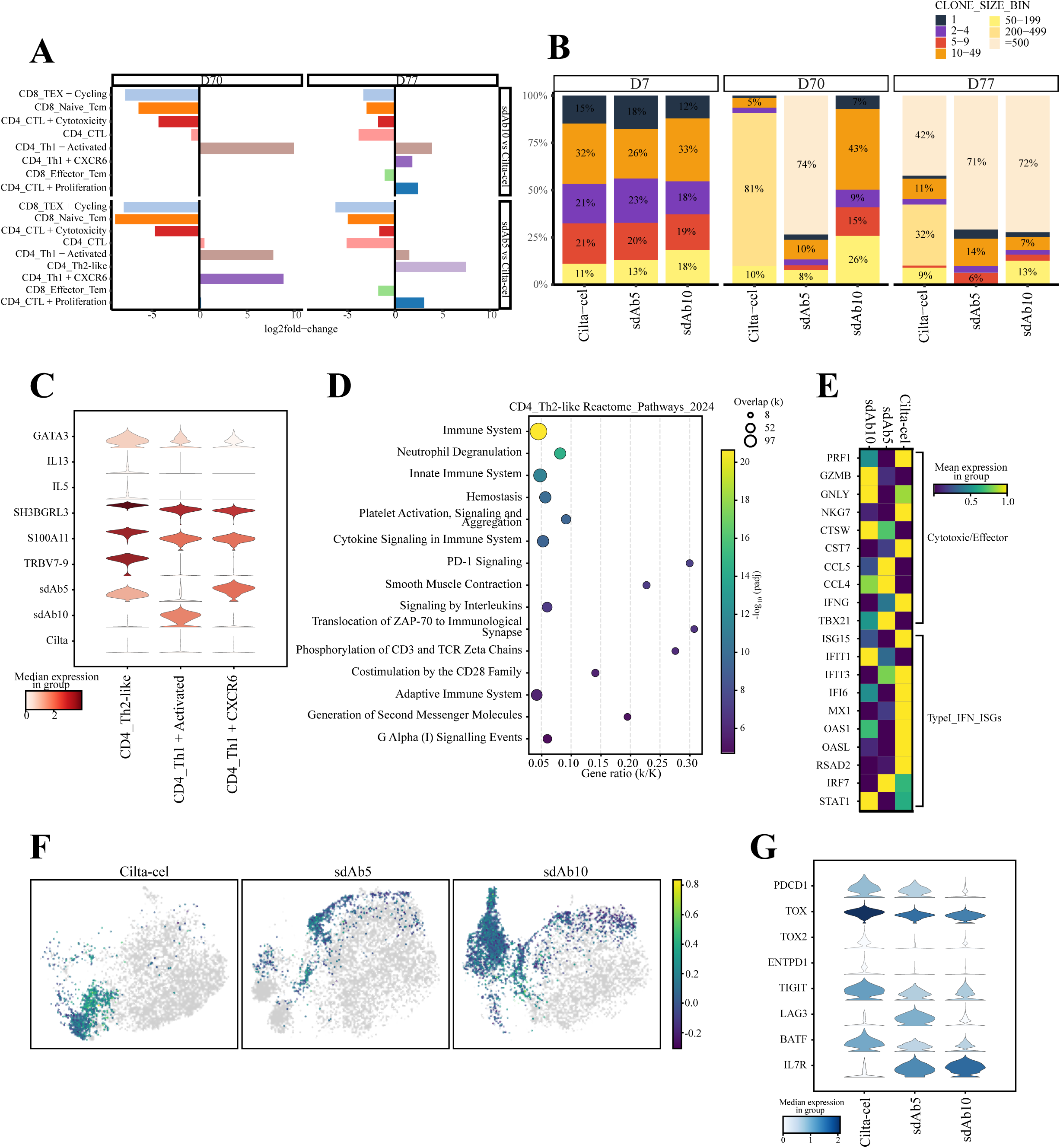
Clonal architecture and transcriptional changes after tumor rechallenge. **(A)** Fold-change in T-cell state frequency across treatments and post-infusion time bins, with negative values indicating enrichment in Cilta-cel and positive values indicating enrichment in sdAb-treated samples. **(B)** Stacked bar plots showing clone-size distribution across treatments within each consolidated sampling window (D7, D70, and D77). **(C)** Violin plots showing expression of selected CD4_Th2-like and related markers across CD4 transcriptional states, including CAR transcripts. **(D)** Reactome pathway enrichment analysis highlighting biological programs associated with the CD4_Th2-like state. **(E)** Heatmap summarizing pseudobulk-derived gene signatures across treatments in CD4_CTL cells, including cytotoxic/effector and type I interferon-associated programs. **(F)** UMAP showing the distribution of CD4 exhaustion scores across treatments in post-rechallenge samples at D77. **(G)** Violin plots showing exhaustion-associated marker expression across treatments at D77.

Collectively, these findings indicate that sdAb-based CAR T cells, and particularly the sdAb5 construct, retain greater transcriptional plasticity during long-term persistence and tumor rechallenge. These cells preferentially occupy transcriptional states associated with migratory potential and functional adaptability and undergo substantial remodeling upon secondary antigen exposure. In contrast, cilta-cel CAR T cells display a comparatively rigid transcriptional landscape characterized by persistent inflammatory activation and limited qualitative reprogramming after rechallenge. These differences in transcriptional flexibility are consistent with the superior tumor control and survival observed in vivo with sdAb-based CAR T cells.

## DISCUSSION

BCMA-directed CAR T-cell therapies have transformed the treatment landscape for patients with R/R MM^1,2^. Nevertheless, a substantial proportion of patients eventually relapse, frequently associated with limited CAR T-cell persistence, progressive functional exhaustion, or reduced efficacy following prior BCMA-targeted therapies. These challenges highlight the need for next-generation CAR designs that preserve long-term functional capacity while maintaining robust antitumor activity. In this study, we demonstrate that single-domain antibody (sdAb)-based CAR recognition domains can shape both antigen engagement and downstream T-cell functional programs, thereby influencing the durability of CAR T-cell responses. Through integrated structural, functional, *in vivo*, and single-cell transcriptomic analyses, we identify sdAb5 as a BCMA binder that supports sustained CAR T-cell activity and preserved responsiveness upon tumor rechallenge.

A key finding of this work is that optimal CAR function is not dictated by maximal antigen affinity alone. Although sdAb10 displayed the highest BCMA binding affinity, sdAb5 consistently mediated superior long-term tumor control and CAR T-cell persistence *in vivo*. These differences are more consistent with a more balanced binding kinetics rather than absolute affinity. Increasing evidence indicates that excessively stable CAR-antigen interactions may impair serial target engagement and promote premature T-cell dysfunction by prolonging signaling events at the immunological synapse^17,25^. Consistent with this concept, the biparatopic sdAb5/sdAb10 construct, despite increased affinity, did not enhance antitumor efficacy. Together, these observations support a model in which kinetic tuning of antigen recognition, rather than maximal binding strength, represents a critical design principle for CAR engineering. Epitope specificity represents an additional dimension of sdAb-based CAR design with potential clinical implications. Our data suggest that sdAb5 recognizes a distinct BCMA epitope from those targeted by ide-cel and cilta-cel. As BCMA-directed therapies are increasingly deployed earlier in the treatment paradigm including antibody-drug conjugates, bispecific antibodies, and CAR T cells^5,6^, therapeutic resistance driven by antigen modulation or epitope masking may become more relevant. In this context, CAR constructs targeting alternative BCMA epitopes may retain activity following prior BCMA-directed therapies. The sustained tumor control and preserved rechallenge responses observed with sdAb5-based CAR T cells are consistent with this possibility and support further exploration of epitope diversification as a strategy to extend the clinical utility of BCMA-targeted immunotherapies. Beyond antigen recognition, sdAb-based CAR constructs displayed distinct functional profiles. *In vitro,* sdAb CAR T cells exhibited minimal tonic signaling and cytokine secretion patterns characterized by preserved IL-2 production alongside restrained inflammatory activation. Excessive early activation and inflammatory signaling have increasingly been linked to CAR T-cell dysfunction and treatment-related toxicities^26^. Although toxicity modeling was not addressed in this study, the restrained inflammatory transcriptional programs observed suggest that sdAb-based CAR designs may support a more balanced activation state, potentially preserving functional capacity while limiting early overstimulation.

Single-cell transcriptomic analyses provided mechanistic insight into these functional differences^20–24^. During the early response phase, all CAR T constructs initiated broadly similar activation programs following antigen engagement. However, cilta-cel displayed stronger induction of inflammatory and interferon-associated transcriptional programs, whereas sdAb-based CAR T cells maintained comparatively restrained activation states and showed a relative enrichment of proliferative CD4 cytotoxic populations. These early differences evolved over time into distinct differentiation trajectories. During long-term persistence, sdAb-based CAR T cells preferentially occupied helper-like transcriptional states associated with functional adaptability, while cilta-cel CAR T cells remained enriched in cycling CD8 populations displaying features consistent with progressive differentiation and exhaustion. Importantly, these divergent trajectories translated into markedly different responses following tumor rechallenge. sdAb-based CAR T cells demonstrated robust transcriptional remodeling and renewed expansion of cytotoxic effector states, indicating preserved functional adaptability upon secondary antigen exposure. In particular, sdAb5 CAR T cells showed a shift towards a CD4 Th2-like transcriptional program characterized by expression of canonical Th2 polarization genes, including GATA3, IL13, and IL5^27–29^. Although the precise functional role of this program warrants further investigation, helper-associated transcriptional states have been linked to enhanced tissue adaptation, immune coordination, and sustained T-cell persistence^28,30,31^. In contrast, cilta-cel showed comparatively limited transcriptional remodeling after rechallenge and maintained strong activation and interferon-associated programs alongside broader induction of exhaustion-associated genes. These findings highlight transcriptional plasticity as a potential determinant of durable CAR T-cell function. The capacity of sdAb5-based CAR T cells to transition between helper, proliferative, and cytotoxic states during long-term persistence may facilitate adaptation to dynamic antigen exposure and tumor evolution. Such flexibility may be particularly relevant in MM, where antigen density, tumor burden, and microenvironmental cues fluctuate during the course of disease.

Several limitations of this study should be acknowledged. The xenograft models used here lack key elements of the human immune microenvironment that may influence CAR T-cell differentiation and persistence^32,33^. Consequently, clinical studies will be necessary to determine whether the functional advantages observed with sdAb-based CAR T cells translate into improved patient outcomes. In addition, while the distinct epitope specificity of sdAb5 may provide advantages in the context of sequential BCMA-directed therapies, the impact of antigen modulation and tumor heterogeneity on this strategy remains to be defined.

In summary, our study identifies sdAb5 as a BCMA-targeting CAR recognition domain that combines optimized binding kinetics, distinct epitope specificity, and favorable transcriptional programming to support durable CAR T-cell function. sdAb5-based CAR T cells maintain restrained inflammatory activation while preserving the transcriptional flexibility required to adapt to repeated antigen exposure. These findings highlight antigen-recognition tuning as a critical determinant of long-term CAR T-cell efficacy, and position sdAb-based CAR architectures as a promising strategy to enhance the durability of BCMA-directed immunotherapies in MM.

## MATERIALS AND METHODS

### Llama immunization and phage display library construction

To generate BCMA-specific single-domain antibodies, a 3-year-old male *Lama glama* was immunized with recombinant human BCMA protein (AcroBiosystems). The immunization protocol consisted of five subcutaneous injections administered in the left subscapular region on days 0, 57, 92, 128, and 214, using incomplete Freund’s adjuvant (Sigma-Aldrich) as emulsifier. Peripheral blood was collected seven days after the final immunization, and peripheral blood mononuclear cells were isolated for total RNA extraction and cDNA synthesis as described^34^. An sdAb phage display library was generated using the pComb3XSS phagemid vector, and *Escherichia coli* TG1 and ER2738 strains were used for library amplification as previously described^34^.

### Phage display selection and screening

The sdAb phage library was subjected to four consecutive rounds of panning against immobilized hBCMA in high-binding 96-well plates (Corning) as described^34^. Antigen concentration was maintained at 5 μg/mL during rounds 1 and 2 and reduced to 2.5 μg/mL for rounds 3 and 4 to increase selection stringency, in combination with progressively more stringent washing conditions. Bound phages were eluted using trypsin (10 mg/mL), and phage titers were determined by infection of *E. coli* ER2738. Phage amplification was performed following infection with M13KO7 helper phage, and phage particles were concentrated and purified by polyethylene glycol-NaCl precipitation. Individual clones derived from rounds 3 and 4 were screened by ELISA for binding to hBCMA using an anti-HA horseradish peroxidase–conjugated antibody (Sigma-Aldrich), and clones displaying cross-reactivity against blocking proteins, including bovine serum albumin and hG9A, were excluded from further analysis.

### Phylogenetic analysis and sdAb selection

Positive phage clones were sequenced by Sanger sequencing, and multiple sequence alignment was performed using Clustal Omega. A phylogenetic tree was generated based on the BLOSUM62 distance matrix^35^, and an average distance threshold of 2.5 was applied to define distinct sdAb families. Based on sequence diversity and binding behavior, five unique sdAbs (sdAb5, sdAb10, sdAb15, sdAb18, and sdAb22) were selected for downstream biochemical, biophysical, and functional characterization.

### sdAb expression and purification

Selected sdAb genes were cloned into the pET28Mod expression vector, a modified pET28a(+) plasmid encoding an IPTG-inducible T7 promoter, an N-terminal OmpA periplasmic localization sequence, and a C-terminal 6xHis-HA tag. Recombinant sdAbs were expressed in *E. coli* BL21(DE3) under optimized induction conditions^34^. Bacterial pellets were lysed either in RIPA buffer (Sigma-Aldrich) or by pressure homogenization. sdAbs were purified by fast protein liquid chromatography, with an initial immobilized metal affinity chromatography step using HiTrap Chelating HP columns (Cytiva), followed by ion exchange chromatography on Q⁺ or S⁻ columns selected according to the predicted isoelectric point of each sdAb. Purified fractions were dialyzed against phosphate-buffered saline and stored at –80°C until use.

### Binding and affinity measurements

Binding activity was initially assessed by ELISA using Greiner Bio-One 96-well plates (Fisher Scientific) coated with hBCMA at 2.5 μg/mL. Bound sdAbs were detected using anti-HA-HRP, and absorbance was measured at 450 nm using a SPECTROstar Nano plate reader (BMG Labtech). EC_50_ values were calculated by nonlinear regression analysis using GraphPad Prism. Bio-layer interferometry (BLI) was performed on an Octet RED96e instrument (Sartorius) using SAX2 biosensors (Sartorius) with immobilization of His-tagged hBCMA. Competitive binding assays were conducted by sequential loading of sdAbs onto antigen-coated biosensors to assess competition for BCMA binding. Surface plasmon resonance (SPR) experiments were performed using a Biacore T200 system (Cytiva) equipped with NTA sensor chips (Cytiva), and binding kinetics were analyzed using either a 1:1 Langmuir or a heterogeneous ligand model, as appropriate.

### Competitive binding analysis of CAR binding domains

Comparative binding analyses were performed by bio-layer interferometry. sdAbs were compared with the scFv derived from humanized version of ide-cel (C11D5.3h) and with sdAbs derived from cilta-cel. HISK biosensors (Sartorius) were used to assess direct binding interactions.

### CAR construct design and lentiviral production

sdAb sequences were cloned into a previously described second-generation pCCL lentiviral vector encoding a CD8α hinge region, a 4-1BB costimulatory domain, and a CD3ζ activation domain under the control of the EF1α promoter. Lentiviral particles were produced in HEK293T cells as described^36^ by transient transfection using pMDLg/pRRE, pRSVRev, and pMD2.G helper plasmids, and viral supernatants were collected for subsequent T-cell transduction.

### *In vitro* characterization of CAR T cells

CAR T cells were generated as described^36,37^. Briefly, primary CD4⁺ and CD8⁺ T cells were activated with anti-CD3/CD28 beads, transduced with lentiviral vectors, and expanded in the presence of interleukin-7 and interleukin-15. CAR T cell function was evaluated using Jurkat transcriptional reporter assays measuring NFAT-GFP, NFκB-BFP, and AP1-mCherry activation^38^. Cytotoxicity assays against MM1S luciferase-expressing target cells were performed as described^34,36,39^, at effector-to-target ratios of 5:1, 2.5:1, 1:1, and 0.5:1 using the Bright-Glo Luciferase Assay. FACS analysis of CD4/CD8 composition and exhaustion markers, including PD-1, TIM-3, and LAG-3, were performed as described^36^. Multiplex cytokine profiling was performed using a 15-plex ProcartaPlex assay according to manufacturer’s instructions.

### *In vivo* tumor models and rechallenge experiments

NSG mice were intravenously inoculated with 3×10^6^ MM1S cells as previously described^36,39^. Fourteen days after tumor infusion, mice received 1×10^5^ CAR T cells by intravenous injection. Tumor burden was monitored longitudinally by IVIS bioluminescence imaging, and survival was analyzed using the Mantel-Cox test. Comparative analyses included sdAb-based CAR T cells, a construct incorporating the C11D5.3h scFv (ide-cel), and a dual sdAb CAR T construct (cilta-cel). To evaluate CAR T cell persistance, mice achieving prolonged disease control were rechallenged with MM1S cells 70 days after initial treatment, and tumor burden was assessed by bioluminescence imaging at days 7 and 15 following rechallenge. CAR T cell persistence in bone marrow and spleen was evaluated by flow cytometry of CD3^+^CD45^+^EGFR^+^ T cells.

### Single-cell RNA sequencing and TCR repertoire analysis

Single-cell RNA sequencing was performed on fluorescence-activated cell-sorted EGFR^+^ CAR T cells isolated from murine bone marrow using the Chromium Single Cell 5’ Reagent Kit (10x Genomics) according to the manufacturer’s instructions. Cells were sorted into phosphate-buffered saline supplemented with 0.04% bovine serum albumin, and cell concentration and viability were assessed using a Nexcelom Cellometer K2. Cell suspensions were adjusted to 700-1200 cells/μL and loaded onto Chromium Single Cell A Chips for gel bead-in-emulsion generation, followed by cell lysis, reverse transcription, barcoding, and cDNA amplification. cDNA quality and quantity were assessed using the Qubit dsDNA HS Assay Kit and the Agilent HS D5000 ScreenTape Assay. Gene expression libraries were generated from 50 ng of double-stranded cDNA and sequenced on a NextSeq 500 system (Illumina; Read 1, 26 cycles; Read 2, 57 cycles; i7 index, 8 cycles) at an average depth of at least 3×10^4^ reads per cell.

T-cell receptor α/β repertoire profiling was performed using the 10x Genomics Single Cell V(D)J Immune Profiling kit. V(D)J transcripts were enriched from barcoded cDNA by targeted amplification and sequenced on an Illumina HiSeq X platform (150-bp paired-end reads; i7 index, 8 cycles) at an average depth of approximately 5×10^3^ reads per cell. Sequencing data were processed using Cell Ranger (v6.0.1) for alignment to the human reference genome (GRCh38) and generation of feature-barcode matrices.

Downstream analyses were performed using Seurat^40^ (v3.1.5), including quality control filtering based on gene counts, UMI counts, and mitochondrial gene content, normalization, identification of highly variable genes, and dataset integration using canonical correlation analysis. Unsupervised clustering was performed at a resolution of 0.8, and dimensionality reduction was carried out using t-distributed stochastic neighbor embedding and uniform manifold approximation and projection. Cell clusters were annotated by manual inspection of differentially expressed genes using canonical marker genes as reference, and paired TCR clonotypes were reconstructed using Cell Ranger (v6.0.2).

### Single-cell gene expression (GEX) and V(D)J processing

GEX and V(D)J libraries were sequenced to a minimum depth of 10,000 and 2,500 reads per cell, respectively, aligned to the human reference genome (GRCh38), and processed with 10x Genomics Cell Ranger (v9.0.0). Downstream analyses were performed using Scanpy^41^ (v1.10.4) and Seurat^42^ (v5.1.0). Cell hashing demultiplexing was carried out with HashSolo^43^ (Scanpy external), using priors (0.05, 0.75, 0.19), and only singlets were retained. Doublets were additionally predicted with Scrublet^44^ and removed. Cells were filtered based on QC metrics computed per cell: cells with ≤200 detected genes (n_genes_by_counts) or ≤500 total UMI counts (total_counts) were removed; cells with mitochondrial RNA fraction ≥10% (pct_counts_mt) were removed; and extreme high-depth/high-complexity outliers were excluded using dataset-specific upper bounds defined as the 99th percentile of detected genes and total UMIs, respectively. Novelty and high ribomosal content cells were flagged. Cells with detectable CD3 expression were retained yielding a total of 24,488 cells. Batch integration across samples was performed with scVI^45–47^, modeling batch as a categorical covariate with default parameters. The learned embeddings were used for downstream dimensionality reduction (UMAP) and unsupervised clustering (resolution 0.3), and clusters were annotated using canonical marker genes as references. TCR analyses were performed using Scirpy^48^ and later scRepertoire^49^.

## ACKNOWLEDGEMENTS

We particularly acknowledge the patients and healthy donors for their participation in this study, and the Biobank of the University of Navarra for its collaboration.

## AUTHORSHIP CONTRIBUTIONS

J.R.R.-M., A.P-L. and F.P. designed the study and funding acquisition. V.I-G. and M.E.C-C. performed the data analysis. V.I-G., F.R., I.A., S.R-D., R.M-T., I.I-S., E.M., F.B-B., P.R-M., L.P-C. and N.G-C. performed wet lab experiments. P.S.M-U., and P.A-R. performed single cell library preparation and sequencing. S.R-D., R.M-T., E.M. and E.I. provided technical assistance. A.A-P., E.T., P.R-O. and J.S-M. provided clinical advice. J.J.L and M.H. contributed to investigation. J.R.R.-M., A.P-L. and F.P. were responsible for research supervision, coordination and strategy. V.I-G., M.E.C-C. and J.R.R.-M. drafted the manuscript. J.R.R.-M., A.P-L. and F.P. reviewed and edited the manuscript. All authors reviewed and approved the final version of the manuscript.

## DATA AND MATERIALS AVAILABILITY

All data needed to evaluate the conclusions in the paper are present in the paper and/or the Supplementary Materials. The scRNA-seq data generated in this study have been deposited in the GEO database (XXX).

