## Supplemental_Material for "Rational engineering of sdAb-based CAR T cells targeting BCMA enhances antitumor efficacy and persistence in Multiple Myeloma"

SUPPLEMENTAL FIGURES

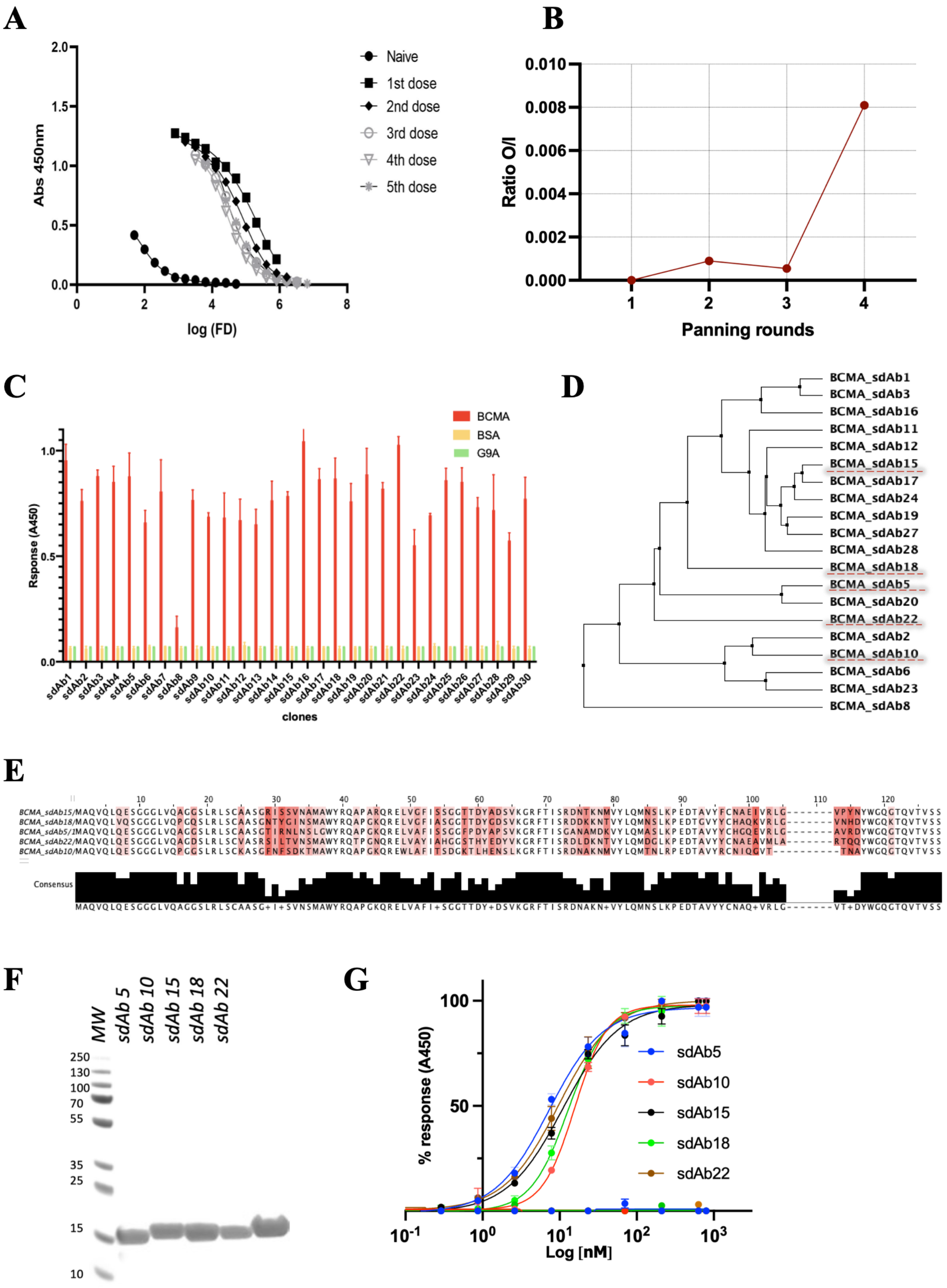


**Figure S1. Identification and Characterization of BCMA-specific sdAbs. (A)** ELISA titration curves demonstrating the BCMA-specific humoral response in llama plasma across different immunization stages. **(B)** Enrichment of BCMA-specific clones represented by the output/input (O/I) phage ratio over successive panning rounds. **(C)** Cross-reactivity test (ELISA) evaluating the specificity of the five candidate sdAbs against unrelated antigens from the same library. **(D)** A phylogenetic tree based on sequence clustering of selected sdAbs, showing distinct families. **(E)** Sequence alignment of the five selected sdAbs, showing conserved framework and divergent CDR regions. **(F)** SDS-PAGE analysis confirming the purity of purified sdAb clones. **(G)** ELISA-based binding curves of sdAbs against recombinant BCMA protein, with $EC_{50}$ values calculated by nonlinear regression.

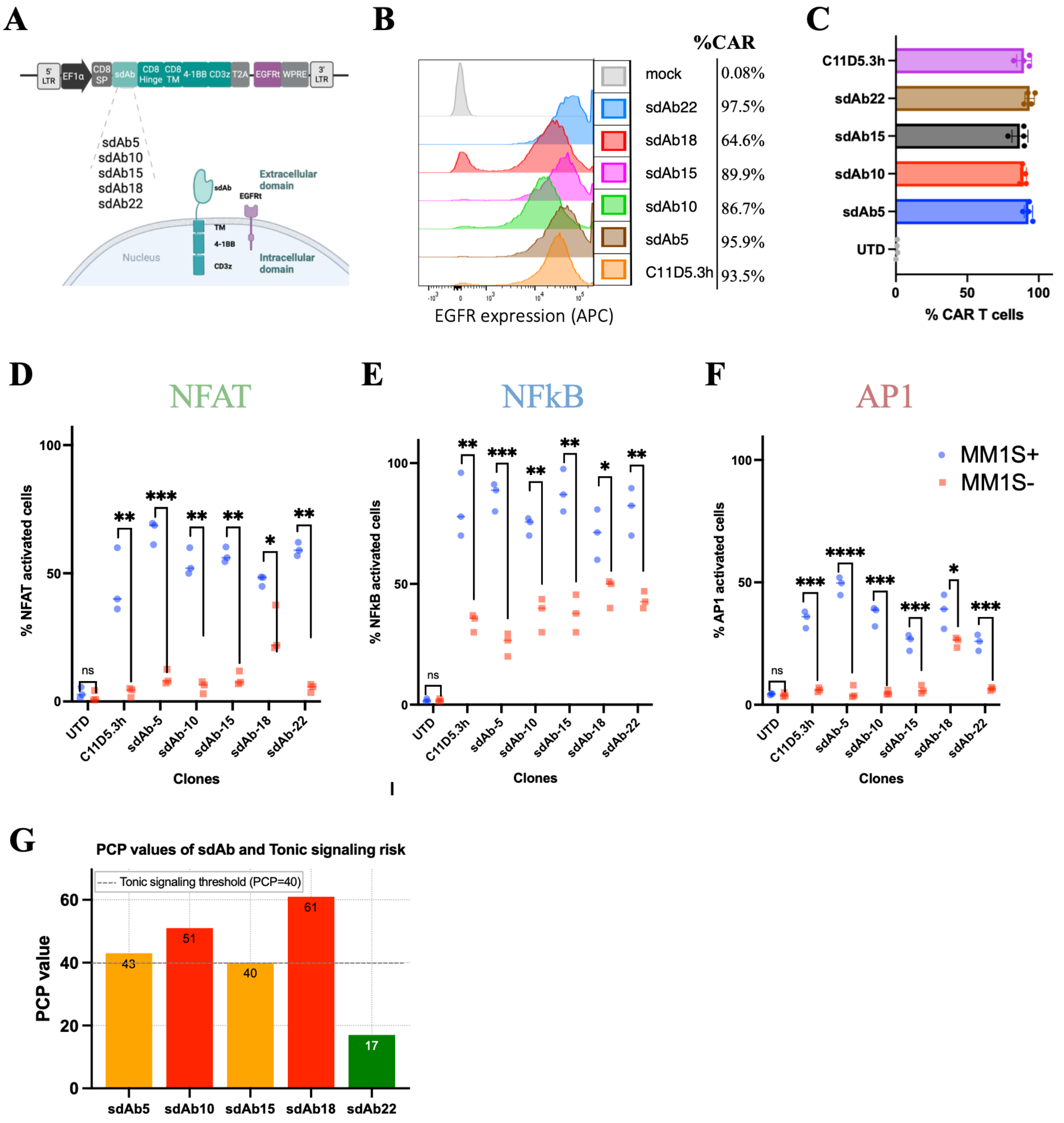


**Figure S2. Transcriptional reporter activation and tonic signaling risk across sdAb-based CAR constructs. (A)** Schematic representation of second-generation CARs incorporating sdAbs, a CD8α hinge, 4-1BB costimulatory domain, and CD3ζ signaling domain. **(B)** Representative EGFR histograms showing CAR surface expression in Jurkat TPR cells. **(C)** Quantification of CAR expression based on EGFR mean fluorescence intensity (MFI) from four independent samples. **(D-F)** Functional activation of NFAT (D), NFκB (E) and AP1 (F) transcriptional reporters in untransduced (UTD) or CAR-expressing Jurkat TPR cells. Cells were analyzed either unstimulated (MM1S$^{-}$) or after 1:1 coculture with MM1S target cells (MM1S$^{+}$). Each dot represents an independent replicate. **(G)** Positively charged patch (PCP) values for each sdAb clone. Dashed line indicates the threshold (PCP = 40) below which tonic signaling is predicted to be minimal. Statistical analysis was performed using two-way ANOVA with Šidák's correction. *P < .05, **P < .01, ***P < .001, ****P < .0001; ns, not significant.

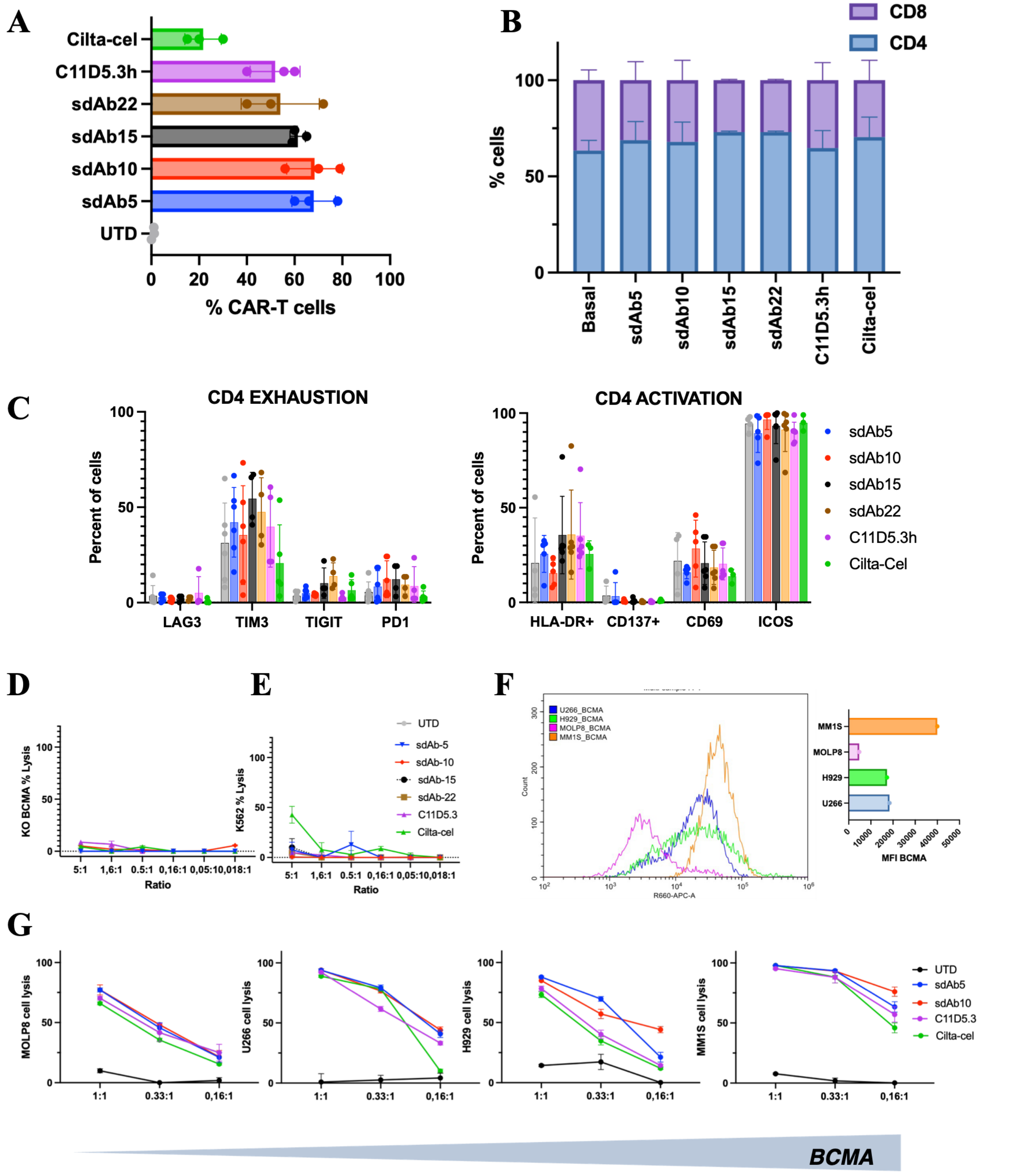


**Figure S3.** ***In vitro*** **phenotypic and functional characterization of sdAb-based CAR T cells. (A)** Transduction efficiency across CAR constructs measured by EGFRt surface expression in T cells from three independent donors. **(B)** Proportional distribution of $CD4^+$ and $CD8^+$ subsets in each CAR T cell product. **(C)** Expression of exhaustion (PD-1, TIM-3, TIGIT, LAG-3) and activation (CD137, CD69, ICOS, HLA-DR) markers on $CD4^+$ CAR T cells following co-culture with $BCMA^+$ target cell lines. **(D)** Cytotoxicity of sdAb-based CAR T cells against MM1S cells lacking BCMA expression (BCMA-KO)

confirms antigen specificity. **(E)** Absence of off-target cytotoxicity against BCMA⁻ K562 cells. **(F)** Surface BCMA expression across MM1S, U266, H929, and MOLP8 cells measured by flow cytometry. **(G)** Antigen-specific cytotoxicity of CAR T cells against multiple myeloma cell lines, demonstrating correlation with BCMA surface density. Data represent mean ± SEM from technical replicates using luciferase-based killing assays at indicated effector-to-target ratios.

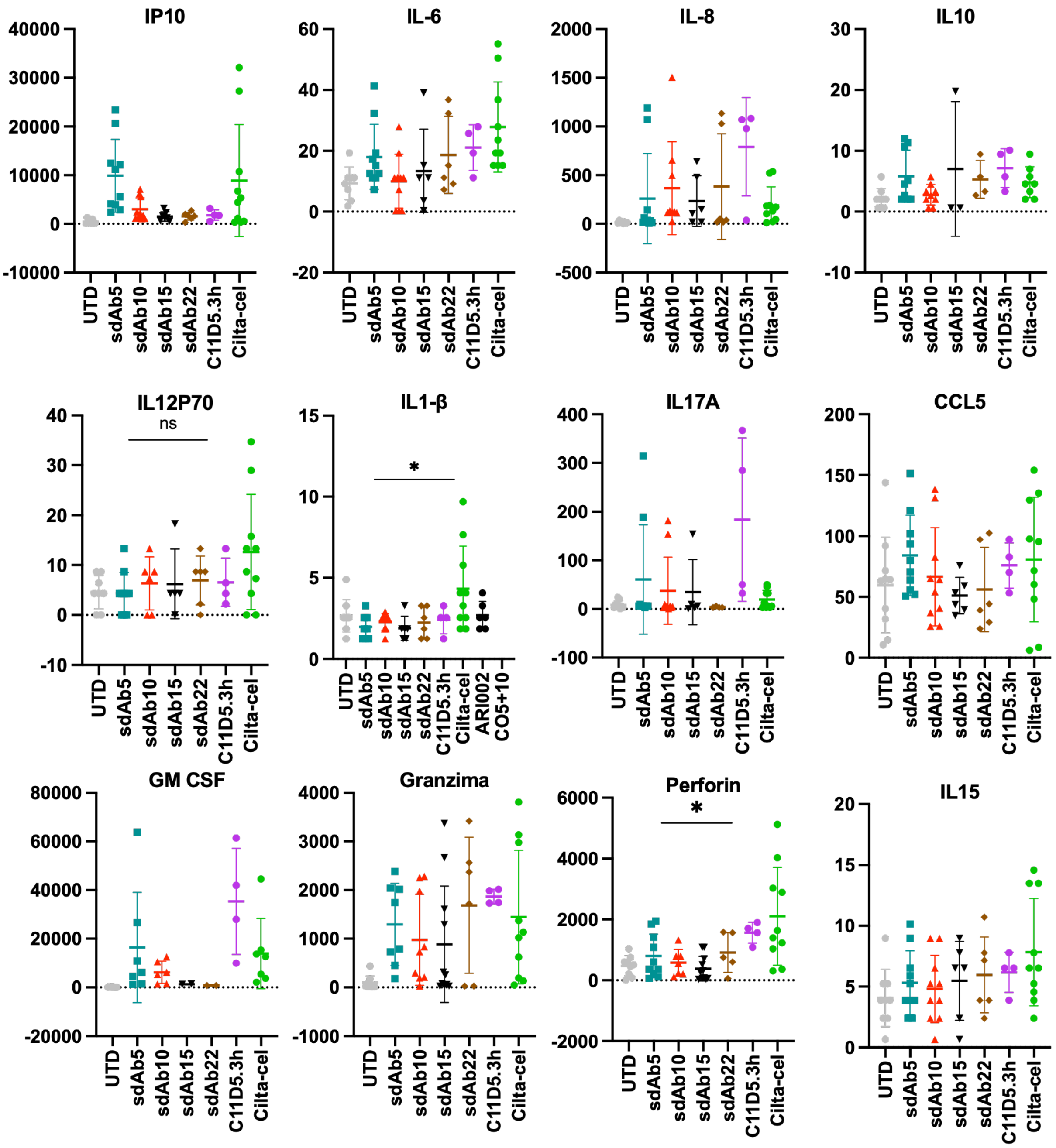


**Figure S4. *In vitro* quantification of cytokine production by sdAb-based CAR-T cells**. Expression levels of a panel of cytokines were measured by Luminex in supernatants from co-cultures with tumoral cells using the indicated anti-BCMA CAR-T cell constructs. UTD cells were used as control. $p < 0.05$

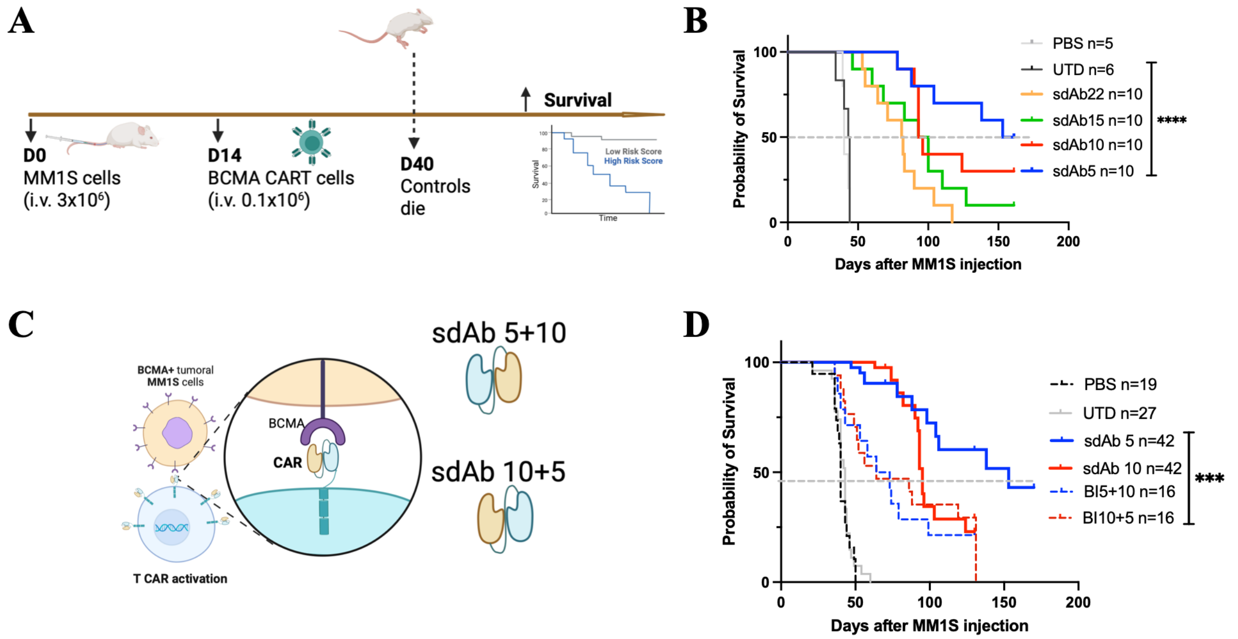


**Figure S5. *In vivo* efficacy of monospecific and biparatopic sdAb-based CAR T cells in a MM1S xenograft model. (A)** Experimental timeline of tumor inoculation, CAR T cell infusion, and bioluminescence monitoring. NSG mice were injected intravenously with 3 × $10^6$ MM1S-$Luc^+$ cells (day 0) and treated with 1 × $10^5$ CAR T cells on day 14. Tumor burden was assessed by IVIS imaging. **(B)** Kaplan–Meier survival curves of mice treated with PBS (n = 5), untransduced T cells (UTD, n = 5), or monospecific sdAb-based CAR T cells (n = 10 per group). **(C)** Schematic of biparatopic CAR constructs incorporating sdAb5 and sdAb10 in tandem, tested in both orientations (5+10 or 10+5). **(D)** Survival curves of mice treated with monospecific CARs (sdAb5, sdAb10), biparatopic CARs, or Cilta-cel (n = 10 per group). Statistical comparisons were performed using Mantel–Cox test. **P < .01, ***P < .001.

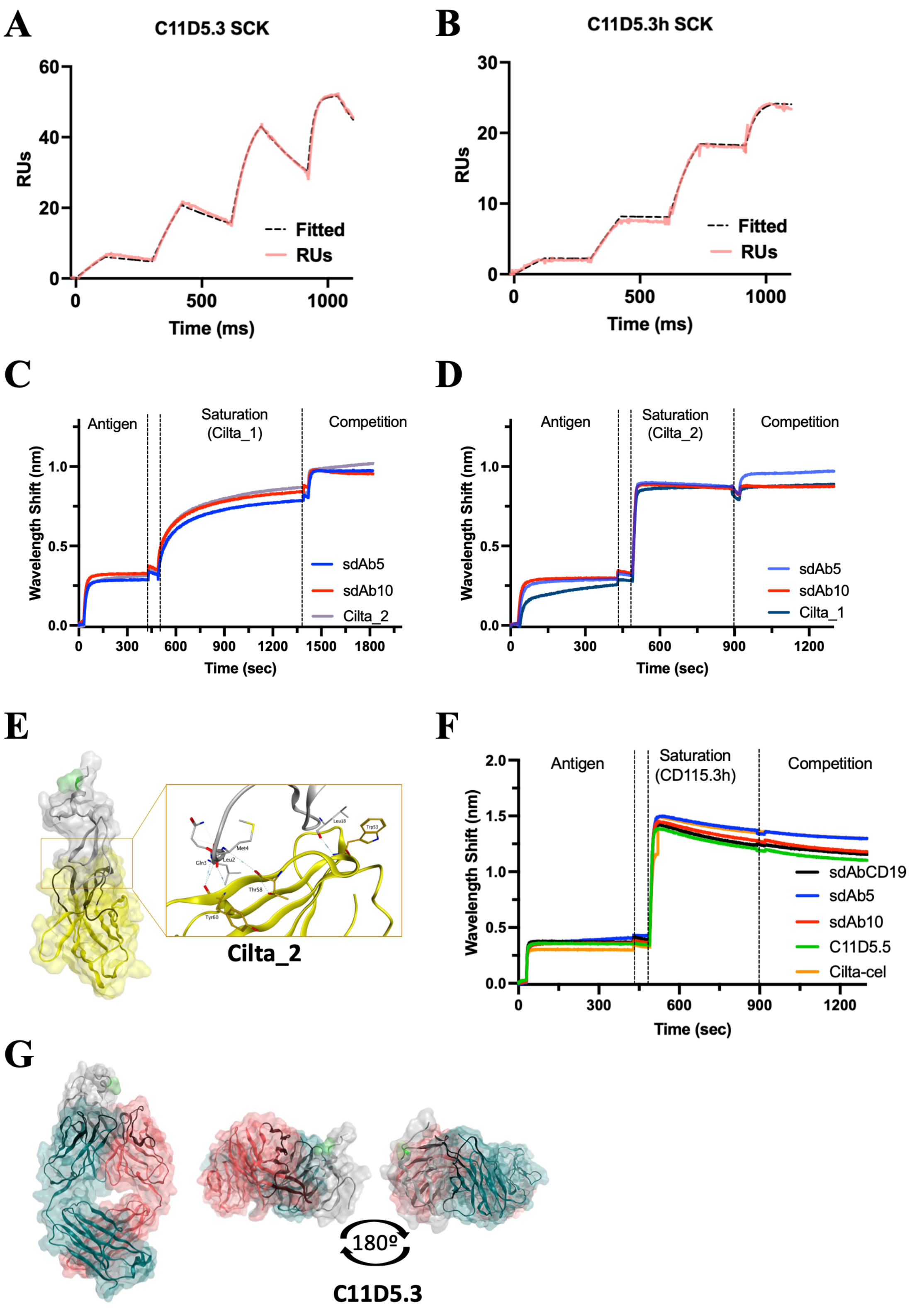


**Figure S6. Epitope mapping of sdAb-based binders by BLI competition and structural modeling. (A-B)** Single-cycle kinetic (SCK) SPR measurements for the C11D5.3 scFv (A) and its humanized variant C11D5.3h (B). **(C-D)** Bio-layer interferometry (BLI) competition assay using Cilta_1 (C) or Cilta_2 (D)

for antigen saturation, followed by addition of sdAb5, sdAb10. **(E)** Structural docking model of sdAb2 from Cilta-cel bound to BCMA, illustrating its epitope location and orientation. **(F)** BLI competition assay in which BCMA was first saturated with C11D5.3h, followed by addition of sdAb5, sdAb10, C11D5.3 or Cilta-cel. The absence of binder displacement indicates non-overlapping epitopes. **(G)** Structural model of BCMA in complex with scFv C11D5.3, showing extended binding interface that overlaps with sdAb5/10 docking regions.

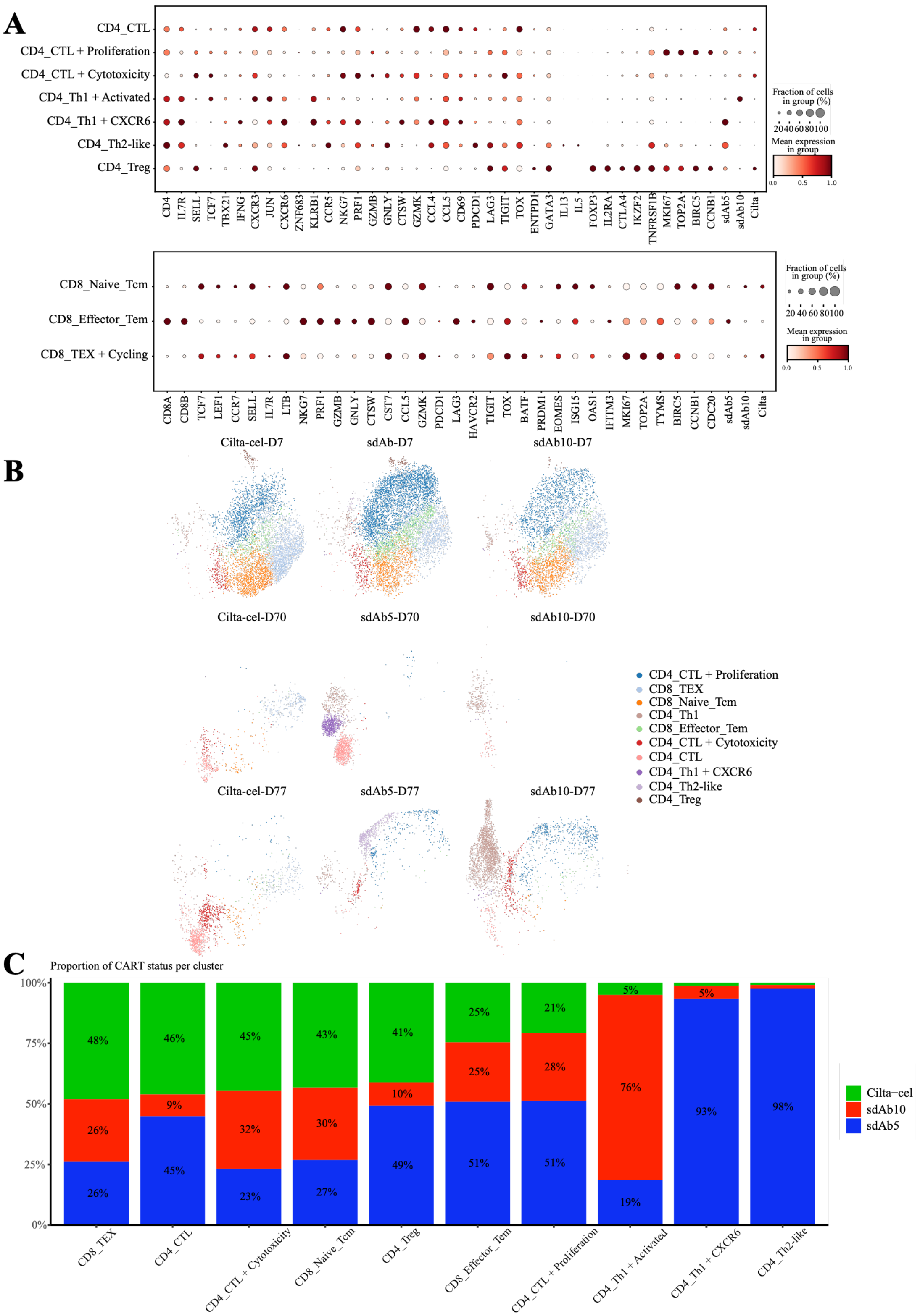


**Figure S7. Annotation and transcriptomic characterization of sdAb-based CAR-T cells. (A)** Supplementary dotplots showing lineage- and state-associated marker genes across CD4 and CD8 compartments. **(B)** UMAPs faceted by day and treatment, colored by the 10 manually annotated T-cell states. **(C)** Stacked bar plots showing the distribution of CAR T-cell products across transcriptional states.

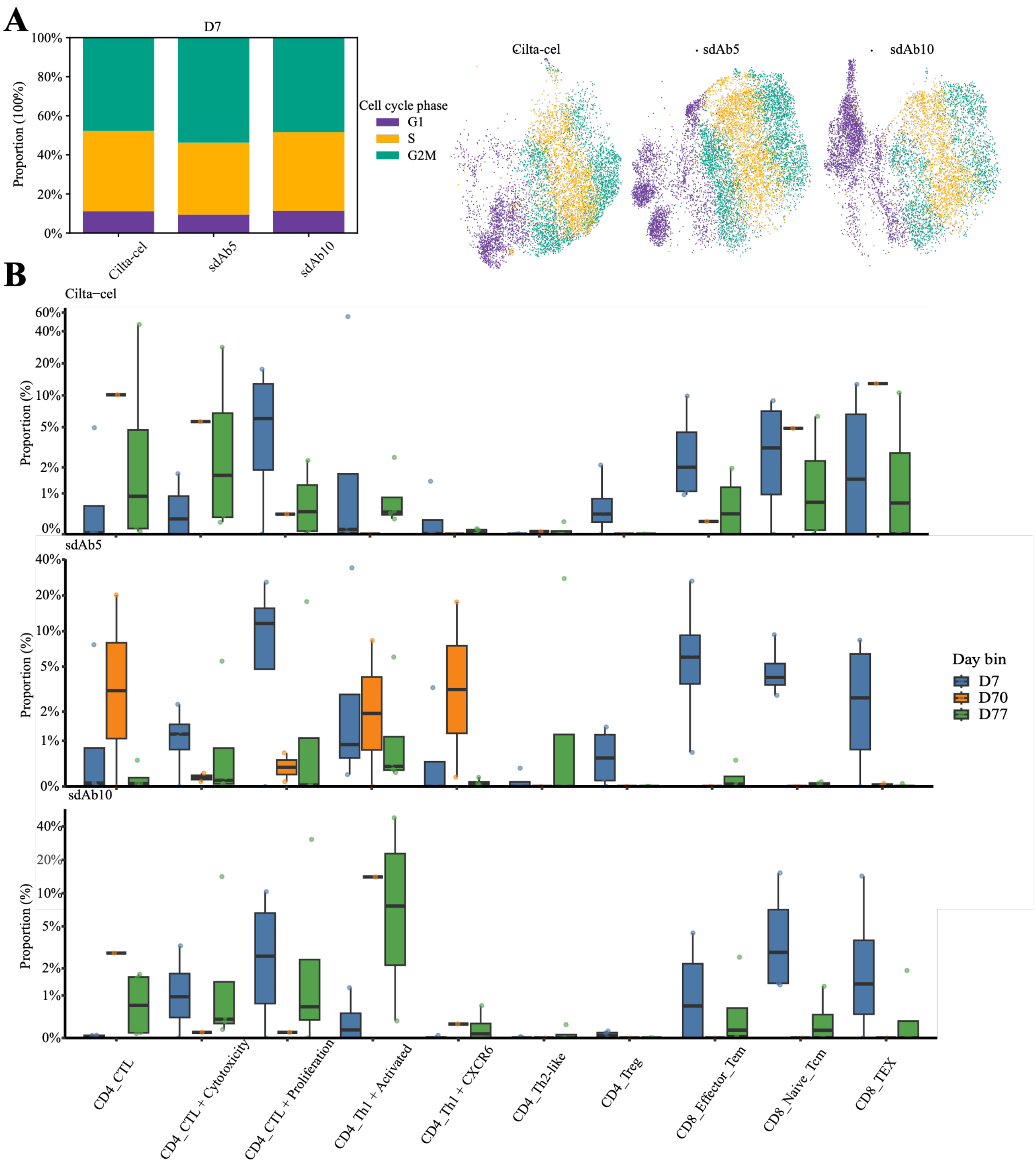


**Figure S8. sdAb-based CAR-T cell compositional variation. (A)** Visualization of cell-cycle phase across treatments at D7, shown together with overall phase proportions. **(B)** Sample-level boxplots showing the distribution of cell-state proportions across treatments and sampling windows. The y-axis is displayed using a pseudo-logarithmic transformation (pseudo-log; sigma = 0.005) to expand the range near zero and improve visualization of low-abundance proportions while retaining 0%.

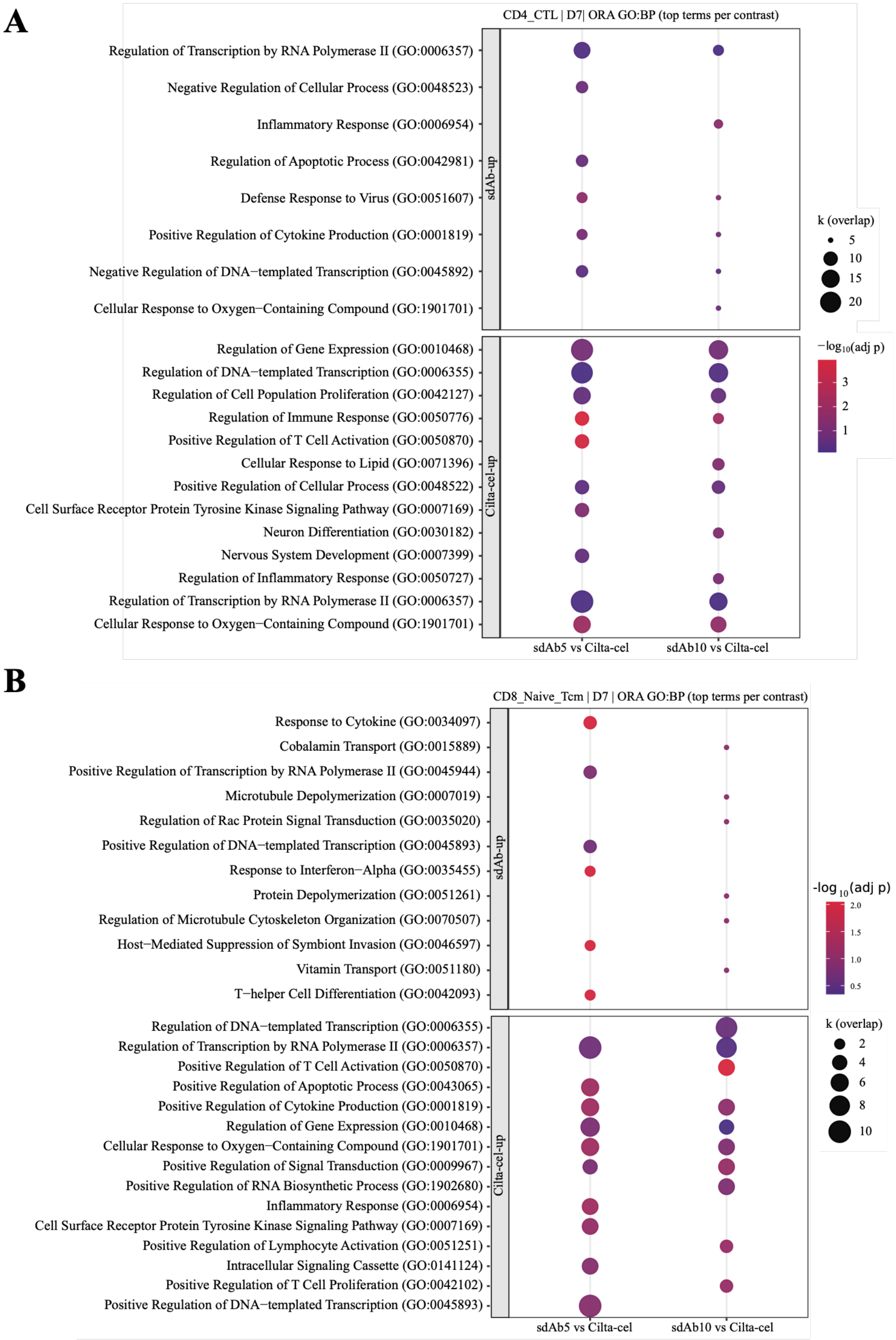


**Figure S9. sdAb-based CAR-T cell functional programs. (A)** GO biological process enrichment analysis for CD8_Naive_Tcm cells at D7. **(B)** GO biological process enrichment analysis for CD4_CTL + Proliferation cells at D7.

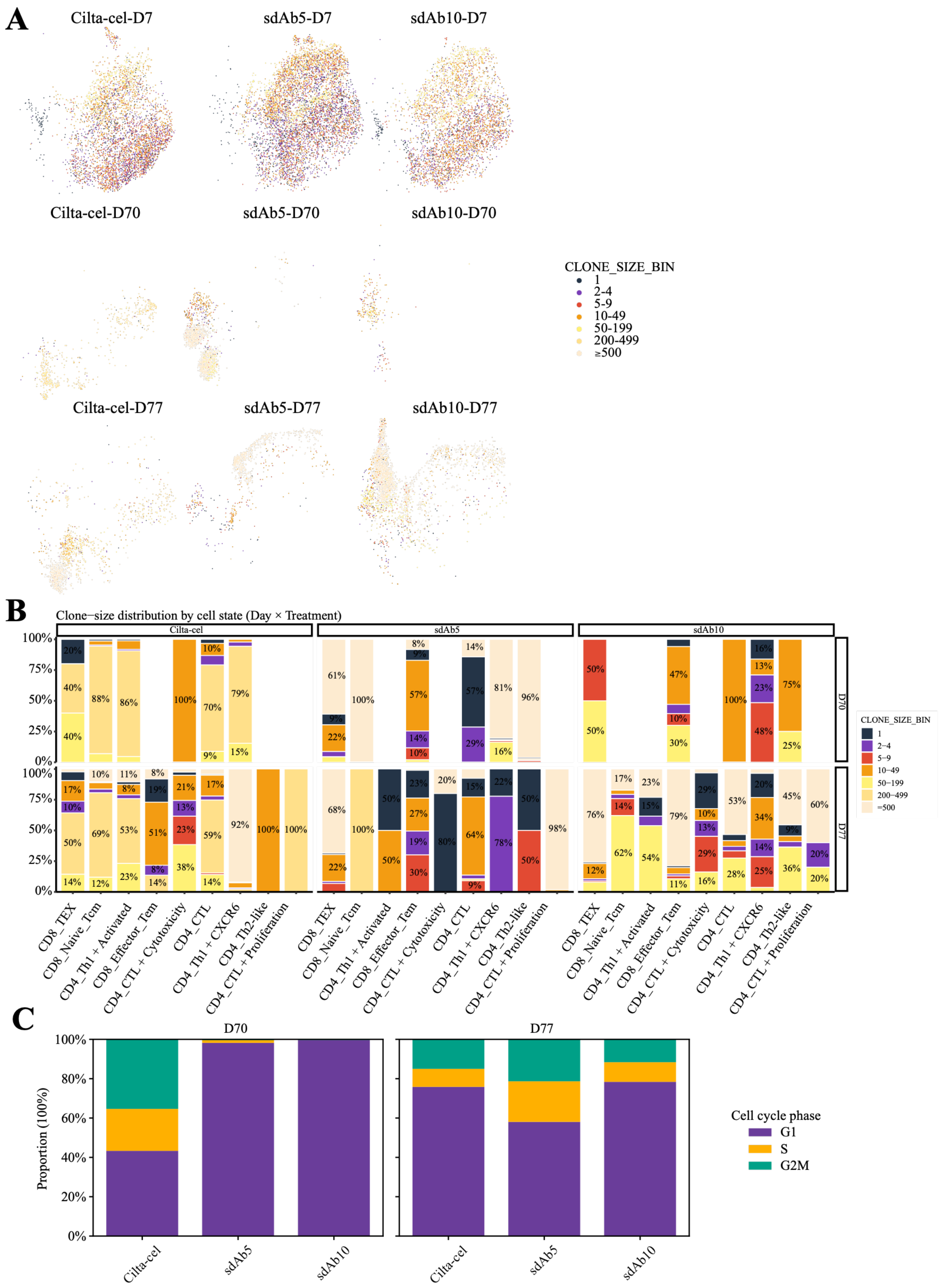


**Figure S10. Clonal analyses of sdAb-based CAR-T cells. (A)** UMAPs faceted by day and treatment, colored by clone-size bin. **(B)** Extended stacked bar plots showing clone-size distribution across T-cell states, treatments, and sampling windows. **(C)** Stacked bar plots summarizing cell-cycle phase proportions across treatments at D70 and D77.

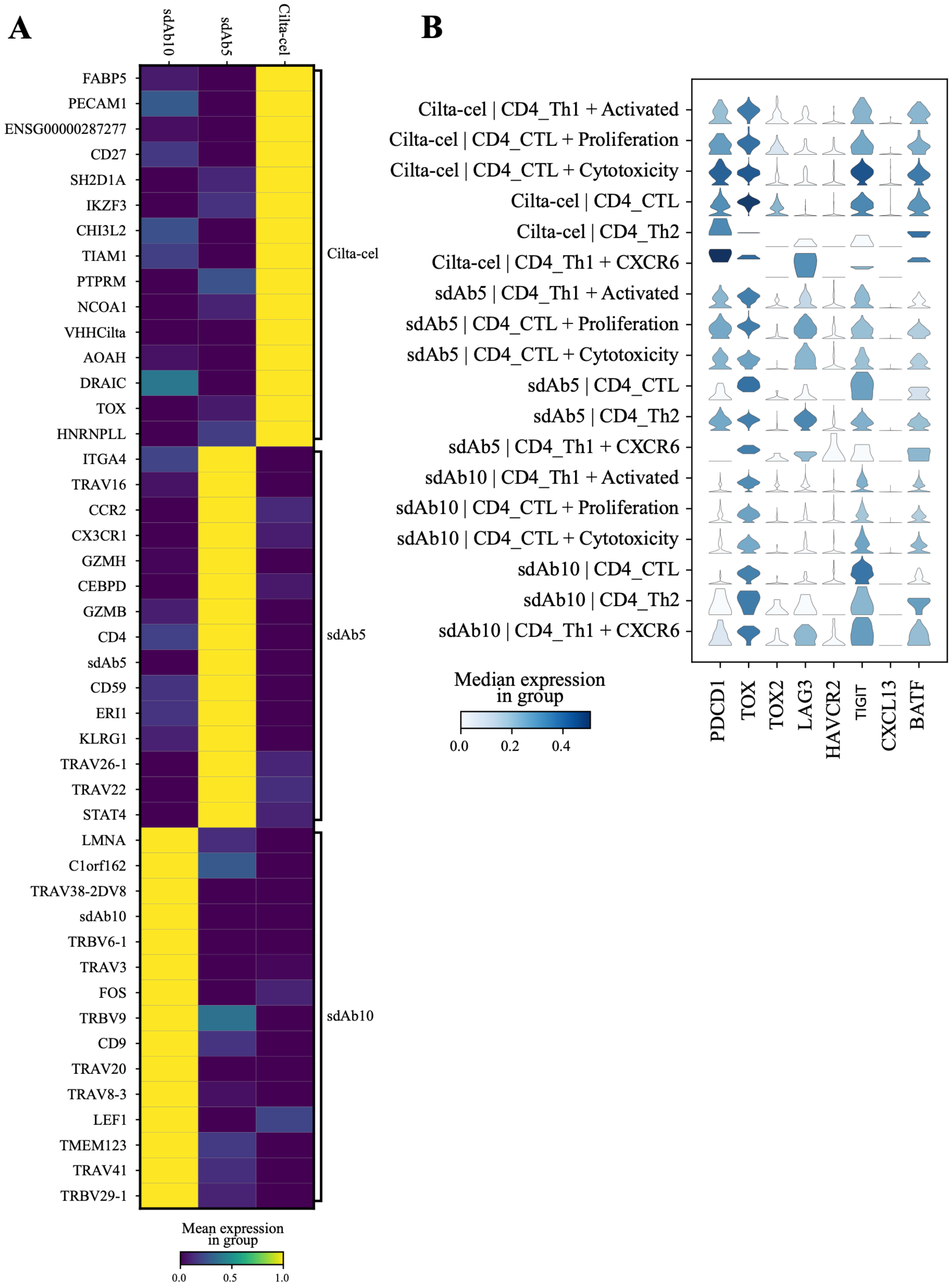


**Figure S11. Functional analyses of sdAb-based CAR-T cells. (A)** Heatmap summarizing treatment-associated gene-expression patterns across post-rechallenge samples. **(B)** Additional violin plots showing exhaustion-associated marker expression across CD4 states and treatments.

**Table S1. Kinetic Parameters of sdAb and CAR-T Binding Domains Against Human BCMA Determined by SPR**

| Antibody | $EC_{50}$(M) | $K_D$(M) | $k_{on}(M^{-1}s^{-1})$ | $k_{off}(s^{-1})$ |
|---|---|---|---|---|
| sdAb5 | 2.66E-09 | 8.62E-10 | 3.07E+06 | 2.40E-03 |
| sdAb10 | 1.99E-09 | 1.75E-10 | 6.88E+06 | 1.07E-03 |
| sdAb15 | 2.65E-08 | 1.16E-07 | 1.17E+05 | 1.36E-02 |
| sdAb18 | 2.17E-08 | 3.92E-09 | 8.58E+03 | 3.36E-04 |
| sdAb22 | 2.02E-08 | 1.38E-09 | 3.97E+06 | 5.48E-03 |
| Cilta_1 | 2.32E-07 | 5.26E-08 | 5.57E+03 | 2.88E-04 |
| Cilta_2 | 3.24E-07 | 1.28E-09 | 1.90E+07 | 2.44E-03 |
| Cilta-cel | 2.64E-08 | 2.79E-10 | 3.15E+05 | 7.64E-05 |
| C11D5.3 | 1.71E-09 | 1.73E-10 | 1.78E+07 | 3.19E-03 |
| C11D5.3h | 2.34E-10 | 1.05E-11 | 6.08E+06 | 6.40E-05 |
